# Nigral-specific increase in ser31 tyrosine hydroxylase phosphorylation offsets dopamine loss and forestalls hypokinesia onset during progressive nigrostriatal neuron loss

**DOI:** 10.1101/2022.11.29.518437

**Authors:** Ella A. Kasanga, Yoonhee Han, Marla K. Shifflet, Walter Navarrete, Robert McManus, Caleb Parry, Arturo Barahona, Vicki A. Nejtek, Jason R. Richardson, Michael F. Salvatore

## Abstract

Mechanisms that augment dopamine (DA) signaling to compensate for tyrosine hydroxylase (TH) loss and delay motor impairment in Parkinson’s disease remain unidentified. The rat nigrostriatal pathway was unilaterally-lesioned by 6-OHDA to determine whether differences in DA content, TH protein, TH phosphorylation, or D_1_ receptor expression in striatum or substantia nigra (SN) aligned with onset of hypokinesia at two time points. At 7 days, DA and TH loss in striatum exceeded 95%, whereas DA was unaffected in SN, despite ∼60% TH loss. At 28 days, hypokinesia was established. At both time points, ser31 TH phosphorylation increased only in SN, corresponding to less DA versus TH loss. ser40 TH phosphorylation was unaffected in striatum or SN. By day 28, D_1_ receptor expression increased only in lesioned SN. These results indicate that increased ser31 TH phosphorylation and D_1_ receptor in the SN, not striatum, augment DA signaling against TH loss to mitigate hypokinesia.

**Highlights:**

–Despite >90% TH and DA loss in striatum, open-field locomotor activity did not decrease
–Early after lesion, DA and TH loss in striatum exceeded 90%. In contrast, DA loss did not occur despite 60% TH loss in substantia nigra (SN).
–TH loss was progressive in the SN, with loss also spreading contralateral to lesion.
–Loss of TH protein in SN preceded cell loss ipsilateral and contralateral to lesion, indicating first stages of nigrostriatal neuron loss begin with loss of TH protein loss.
–TH phosphorylation at ser31 in SN was associated with less, if any, DA loss compared to TH protein loss.
–TH phosphorylation at ser40 did not change in either region and at any time during TH loss, suggesting no contribution to differences in DA loss against TH loss.
–Expression of the D1 receptor increased 2.5-fold in the SN late, but not early, after lesion, suggesting a post-synaptic receptor response to offset DA loss in SN.
–No increases in TH phosphorylation or D1 receptor expression in striatum at any time after lesion induction, indicating that compensatory mechanisms occur only in substantia nigra, but not in striatum, to delay onset of hypokinesia.

## Introduction

Strategies to slow Parkinson’s disease (PD) progression is a major knowledge gap. A critical challenge for firm diagnosis of PD is the need for diagnostic biomarkers and related prodromal signs early in the disease process [1–4]. When motor impairment strikes, major loss (∼80%) of tyrosine hydroxylase (TH) and other dopamine (DA)-regulating proteins has already occurred in nigrostriatal terminals in striatum [5–7]. This metric of striatal TH or DA loss has been supported in non-human primate PD models, as even 60% loss does not promote onset of hypokinesia [7]. The major loss of TH in striatum suggests that presumably little DA-biosynthesis capability remains in striatum after PD diagnosis. However, motor impairments continue to worsen in severity even though DA markers are virtually undetectable 5 years after diagnosis [6]. Paradoxically, restoration of motor capabilities in patients may not be possible with recovery of striatal TH [8]. Conversely, recovery of motor function without commensurate increases in DA or TH in striatum can occur [9–14] With the predominating hypothesis that striatal DA deficits are central to parkinsonian signs like hypokinesia, these ambiguities continue to thwart establishing a cohesive and mechanistically-grounded therapeutic blueprint to promote motor recovery.

Compensatory processes have been long-thought to augment DA signaling in striatum to delay onset of parkinsonian signs [15–20]. The candidate mechanisms mediating these compensatory processes are many, and include increased activity of other basal ganglia circuits, increased metabolic or synaptic activity in motor cortex, cerebellum or thalamic nuclei, and, finally, inter-hemispheric-based interactions [18]. Striatal-based increases in components of DA neurotransmission have been most extensively studied at several steps, including increased DA synthesis, DA metabolism or DA receptor expression. However, well-controlled primate PD studies revealed opposing conclusions as to whether increased striatal DA metabolism was indicative of increased DA signaling to delay parkinsonian sign onset [7, 19]. It is now clear that increased striatal DA metabolism occurs only after motor impairment, regardless of possible recovery [17]. As a result, alternate ompensatory mechanisms in DA signaling have been proposed to exist outside of the striatum [17, 18].

The rate of loss of DA-regulating proteins in human PD proceeds at a slower rate in the nigrostriatal somatodendritic compartment in substantia nigra (SN) compared to striatum [6,7,20–22], suggesting this may be a compensatory mechanism to delay parkinsonian signs. Nigral DA markers remain at 10-50% of normal, long after those in striatum are undetectable in patients [6,20–22]. Nevertheless, the possibility that nigral DA signaling affects locomotor function is rarely considered, and instead, remaining TH+ cells are quantified as a static biomarker of neuron viability. Yet, multiple lines of evidence challenge the perspective that deficiencies in striatal DA signaling are the chief regulating mechanism of parkinsonian sign onset or severity [7, 9,14,18,19,21–27]. Moreover, local modulation of nigral DA signaling affect locomotor activity levels autonomously from any change in striatal DA [9,28,29].

Using the well-established 6-OHDA hemi-lesion rat PD model, we addressed how the biosynthesis and post-synaptic signaling aspects of DA neurotransmission changed in the striatum and SN against the timing of hypokinesia. Changes in TH phosphorylation stoichiometry at ser31 or ser40 during lesion progression were evaluated against of DA vs TH loss as a compensatory mechanism, as suggested by changes in striatal TH activity or phosphorylation in PD models [30–34] and human PD [35]. Finally, as DA D_1_ agonists can mitigate parkinsonism [36], and their function in the SN affects basal ganglia function and locomotor activity [9, 29,37–39], we also evaluated if expression of the DA D_1_ receptor [40–43] changed in response to DA loss. Our results clearly indicate increased TH phosphorylation at ser31 offsets TH protein or cell loss to maintain DA levels only in the SN, and temporally coincided with forestalling hypokinesia. With further nigrostriatal TH protein and cell loss, and eventual onset of nigral DA loss, increased D_1_ receptor expression, specifically in the SN, was engaged. Together, these results clearly point to plasticity in DA signaling in the SN as a mechanism to mitigate hypokinesia against progressive loss of the nigrostriatal pathway in PD.

## Methods

### Study design

The study objective was to determine longitudinal differences in DA regulation following induction of nigrostriatal lesion, with particular attention to unique distinctions in DA signaling between striatum and SN. A total of 65 male Sprague-Dawley rats, age 3 months old, were used between identical studies to obtain different outcomes (neurochemistry for DA tissue content and TH protein and phosphorylation for one study and immunohistochemistry for total neuron and TH-positive neurons for the other study). Rats were placed into a day 7 or day 28 cohort at random, but were then evaluated for pre-lesion locomotor activity levels (see below). A sham-operation group was included as a control for evaluation of potential differences in striatal or nigral DA measures contralateral to lesion and for possible influences of the operation itself. For neurochemistry and immunohistochemistry assessments, rats were subdivided into an *n=*40 or 25, respectively. Based upon our previous work to assess DA loss with this strain and using a 6-OHDA lesion model [31], the power in the current study is 0.73-0.96. In the neurochemistry study, rats that did not meet lesioned criterion by day 7 (see *forepaw adjustment steps* below) continued in the study until completed on day 28, if not designated for dissection in the day 7 cohort. Evaluation of motor function and neurochemistry was done on all rats regardless of FAS outcome on day 7. For statistical analysis in the neurochemistry cohort, rats that did not meet a minimum of >25% FAS were excluded from motor and neurochemistry analysis. A total of 6 rats of 25 6-OHDA-lesioned rats did not meet the FAS threshold (2 in day 7 cohort and 4 in day 28 cohort). In the immunohistochemical study, the criterion for exclusion from statistical analysis was for TH protein loss to not exceed at least 80% loss in lesioned striatum versus levels in striatum contralateral to lesion. A total of 3 rats of 14 did not meet the threshold for >80% striatal TH protein loss.

### Animals

Male Sprague-Dawley rats age 3 months old were obtained in 3 different cohorts from Charles River (Wilmington, MA). They were used in two studies with identical experimental design to evaluate changes in DA and TH regulation and nigrostriatal neuron expression over two time points (7 or 28 days) following induction of nigrostriatal neuron loss by 6-OHDA. One study (DA regulation impact) was designated for analysis of DA tissue content, TH protein, TH phosphorylation at ser31 and ser40, and DA D_1_ receptors by quantitative western blot, and the other study (neuronal impact) was designated for analysis of TH or Nissl cell count number, with accompanying verification of TH loss in the striatum by quantitative western blot. Rats arrived into the vivarium maintained at ambient conditions in a reverse light-dark cycle and fed *ad libitum.* Rats were acclimated for at least one week prior to handling for assessment of pre-nigrostriatal lesion locomotor performance.

### 6-OHDA lesion induction and sham-operation

Rats were deeply anesthetized with 2-3% isoflurane and placed into a stereotaxic device (Kopf). After incision in the scalp, Bregma was identified, and small holes were drilled into the skull at locations to target the medial forebrain bundle (mfb) in both hemispheres. Targeting the mfb ensures loss of TH and DA in both striatum and SN [31]. Using a 26 gauge needle (Plastics One, cat#, Roanoke, VA), freshly-prepared 6-OHDA (16 µg in 4 µg/µl) in 0.02% ascorbic acid in sterile saline was infused into the mfb at coordinates relative to Bregma (in mm) AP −1.8, ML 2.0, DV −8.6 at a rate of 1 µl/min. The needle was left in place for 10 min after infusion prior to withdrawal. The mfb on the side contralateral to lesion received an infusion of the same volume of 0.02% ascorbic acid in sterile saline (without 6-OHDA) to control for impact that infusion or the needle may have on dependent measures in the study.

To determine if unilateral 6-OHDA lesion affected motor or DA-related measures in the hemisphere contralateral to lesion, a sham-operated group was included and used identical time-points post-surgery. The sham-operated group received 0.02% ascorbic acid in sterile saline delivered into the mfb in one hemisphere. The mfb contralateral to the sham-operated side remained intact.

After 6-OHDA lesion induction or sham-operation, the rats were further subdivided into two separate cohorts consisting of a 7-day post-lesion and 28-day post-lesion (or post sham-operation) group. Further locomotor testing ensued on day 7 post-lesion in both groups, with additional locomotor assessments continuing in the 28-day post-lesion group every 7 days (day 14, 21, and 28 post-lesion or sham-operation).

CNS tissues were obtained immediately following the final locomotor assessments in both cohorts. The nucleus accumbens (NAc) and ventral tegmental area (VTA) were also taken with the striatum and SN as a reference to lesion-specific effects in the nigrostriatal pathway. The dissection procedure and ensuing tissue processing for contemporaneous assessment of DA and other monoamines with specific proteins (like TH) and site-specific TH phosphorylation in the same tissue dissection sample has been well-established [9, 31, 44–47].

### Assessment of motor function

#### Forepaw adjustment steps

Rats were acclimatized to the procedure for 3-5 days by daily handling to become familiar with the experimenter’s grip, as well as the required positioning for the test. To begin the test, the rat is held at one end of tape (90 cm long) on a table with hind legs placed in one hand, by holding the rat at a 45° angle facing down so that, with the other hand, the same arm of the rat is held (ex: right hand holds right arm) so that its weight is now only on one forelimb. The paw is then placed in front of the end line of the tape to evaluate the number of steps initiated in 10 seconds while the rat is moved across the length of the tape on the table. This is done for 6 trials per forepaw: 3 forehand trials (lateral steps toward the thumb of the paw that is stepping) and 3 backhand trials (lateral steps towards the pinky of the paw that is stepping. The starting paw is alternated for each animal on consecutive test days. In the 7- and 28-day day cohorts, baseline (pre-lesion or sham-op) measures were obtained from both forelimbs. In the 7-day cohort, one more assessment was taken 7 days after lesion or sham-op. In the 28-day cohort, following 6-OHDA or sham-operation, evaluations took place on a weekly (7 days apart) basis.

Prior to group assignment, we verified no significant differences in in left or right forelimb use, which averaged 93.4 ± 2.3 and 88.6 ± 3.0 and 92.1 ± 2.2 and 88.1 ± 3.4 in the 7-day and 28-day cohorts, for left and right limb respectively (Suppl. Fig. 1). Sham operation had little, if any effect, on forelimb use in the 7-day cohort (t=1.01, ns, df=7). In the 28-day cohort, sham-operation also had minimal effect, with only a 10% increase in forelimb use at 21 days (Suppl. Fig. 2).

#### Open-field locomotor evaluation

Spontaneous locomotor activity was monitored for one hour in a darkened room during the rat awake cycle using automated activity chambers (VersaMax Animal Activity Monitoring System, Columbus Instruments, Inc., Columbus, OH). Activity monitors consist of a 41×41×31 cm^3^ plexiglass box with a grid of infrared beams to automatically record activity. Locomotor activity was evaluated in movement parameters defined as ambulatory time (AT, time of initiated movements), total distance (DT, cm) and speed (cm/sec). Three assessments were taken to establish a baseline for each rat prior to induction of 6-OHDA lesion or the sham-operation. In the 7 day cohort, one more assessment was taken 7 days after lesion or sham-op. In the 28 day cohort, following 6-OHDA or sham-operation, evaluations took place on a weekly (7 days apart) basis.

Rats assigned to the Sham and 6-OHDA groups had no significant difference in locomotor activity prior to treatments (Suppl. Fig. 3). Sham-operation had minimal effect on locomotor activity. Distance travelled was unaffected at any time in the 7-day or 28-day cohort on the 7^th^ day, (Suppl Fig. 4 A,B), and only a slight ∼15% increase in time spent moving at days 14 and 21 in the 28-day cohort (Suppl Fig. 4 C,D). Movement speed was unaffected by the sham-operation in either cohort ((7-day cohort, t=0.24, ns,df=7) (28-day cohort (F_(4,24)_ = 1.07, ns), (data not shown)).

#### Tissue dissection

In the DA regulation impact study, CNS tissues were dissected immediately following locomotor assessments on day 7 in the day 7 cohort or day 28 in the day 28 cohort. Rats were briefly rendered unconscious with isoflurane and immediately decapitated for rapid brain removal. The whole brain was placed into an ice-cold coronal rat brain matrix to dissect dorsal striatum, SN, NAc, and VTA, as described [44]. Tissue dissections were immediately chilled in precooled microfuge tubes on dry ice and stored at −80°C until further processing.

In the immmunohistochemical study, the striatum was dissected fresh frozen when either time point was reached, and the remaining midbrain was dropped fixed into 4% paraformaldehyde for 7 days at 4°C. Cryoprotection consisted of subsequent immersion into 30% sucrose with 0.1% sodium azide. These tissues were processed for immunohistochemistry as described below.

#### Immunohistochemistry

The midbrain was divided into left and right hemispheres prior to sectioning. To match the coronal sections of the left and right hemisphere, the two hemispheres were placed together and snap-frozen in dry ice, then cut together into 30 µm sections using a sliding microtome (Thermo Fisher Scientific, HM450). Each section was separated into left and right hemispheres and placed in individual wells containing cryoprotectant solution (30% glycerol + 30% ethylene glycol + 0.024M phosphate buffer) until staining. Every 6th section was stained for Nissl, and the immediately adjacent sections were stained for TH. For Nissl staining, the sections were mounted on the glass slide and cresyl violet acetate solution (0.5% cresyl violet acetate and 0.3% glacial acetic acid) was used to stain Nissl bodies (rough endoplasmic reticulum) in the cytoplasm of neuronal cells. For TH staining, free-floating sections were washed for one day with phosphate-buffered saline (PBS) at 4°C, then permeabilized with PBS + 1% Triton X-100 for 15 min at room temperature. After quenching of endogenous peroxidase with 2.5% H2O2 in 75% methanol, the sections were incubated for 1 h with a blocking solution (4% goat serum + 5% bovine serum albumin + 0.5% Triton X-100). The sections were incubated for 48 h at 4°C with primary mouse anti-TH antibody (Cat#MAB-318, Millipore Sigma) at 1:1000 dilution. The sections were then incubated for 1 h at room temperature with biotin-conjugated goat anti-mouse secondary antibody (Cat#BA-9200-1.5, Vector Laboratories) at 1:200 dilution. The sections were washed with PBS and incubated with Vectastain Avidin-Biotinylated Peroxidase Complex (ABC) Detection Kit (Cat#PK4000, Vector Laboratories) and 3,3’-diaminobenzidine (DAB) Substrate Kit with Peroxidase and Nickel (Cat#SK4100, Vector Laboratories) for development. The sections were mounted on glass slides and dehydrated in a series of ethanol solution from 30-100% prior to cover slipping.

#### Unbiased stereology

Nissl+ and TH+ cells were counted in sections containing the substantia nigra pars compacta (SNc) using Stereo Investigator software (MBF Bioscience, USA) and an upright digital microscope (Leica DM4 B Automated Upright Microscope System Cat#DM4B, Leica Biosystems, USA). An optical fractionator was chosen as a stereoscopic probe to measure the size of the cell population. The sampling parameter for both cell counts was set with a counting frame of 200 × 200 µm and a dissector height of 10 µm. Grid sizes of 200 × 200 µm and 300 × 300 µm were set for TH+ and Nissl+ cell counting respectively.

To quantify both cell populations, a contour was created around the SNc region according to The Rat Brain in Stereotaxic Coordinates (Paxinos & Watson, Compact 7th edition, Academic press) using the 4X objective and then counting was performed using the 20X objective. The mounted thickness was measured at each sampling site and the average was calculated to be 11.2µm. Cell counts were estimated from 4-6 sections per sample for TH and 3-4 sections per sample for Nissl. The coefficient of error, calculated by the Gundersen formula, was less than 0.1 for both cell counts. Sham-operation had little or negligible impact on TH or total neuron number (Suppl. Fig. 5A, B), with total TH cell number ranging ∼5100-6900 cells, and total neuron numbers ranging from 15,300-17500 cells. Thus, TH+ cells comprised roughly 30-45% of total neurons counted in SN.

### Tissue processing for DA, DOPAC, and NE tissue content and protein expression

Fresh-frozen tissues were sonicated in ice-cold perchloric acid solution [44] and centrifuged to segregate precipitated protein from the supernatant. The supernatant was analyzed in-house by HPLC (Thermofisher) for DA, dihydroxyphenylacetic acid (DOPAC) for assessment of DA turnover, and norepinephrine (NE). A standard curve, ranging from 1.56 to 800 ng/mL was used to quantify DA, DOPAC, and NE. The resulting values were calculated to total recovery in the sample and normalized against total protein recovered.

The precipitated protein was sonicated in 300-400 µl 1% sodium dodecyl sulfate solution [44] and 5 or 10 µl aliquots were assayed for total protein by the bicinchoninic acid method against an albumin standard curve. Sample buffer with dithiothreitol as a reducing agent was then added to the sonicated samples for SDS-PAGE for multiple quantitative Western blot assessments. The samples for all treatments and groups were represented and balanced within each blot, so that individual differences in treatments were captured within each blot.

Tyrosine hydroxylase protein was quantified against a calibrated standard traceable over multiple studies [9,31,44–47]. Optimal total protein load for TH determination in striatum and SN was 4 and 8 µg, respectively, using the primary antibody obtained from Millipore (Temecula, CA, cat #AB152). TH phosphorylation stoichiometry was determined for Ser31 using primary antibody characterized for specificity at 1 µg/ml dilution [44]), and Ser40 using primary antibody from PhosphoSolutions cat # p1580-40 at 1:1000 dilution). After determining total TH content, a nominal total TH load of 1 or 2 ng was assayed for TH PS using a calibrated standard for each phosphorylation site ranging from 0.01 to 0.60 ng of phosphorylated ser31 and 0.002 to 0.02 ng of phosphorylated ser40 [44, 45].

The DA D_1_ receptor exists in a glycosylated and non-glycosylated form in the rat basal ganglia at 75 and 50 kDa [41, 42]. The expression of the receptor was quantified at both molecular weight forms using anti-DRD1 primary (Sigma Aldrich, cat#D2944) at 1:1000 dilution. Prior for use in western blotting, the primary was preincubated for 1 hr at RT with 1 µg/ml AffiniPure Fab Fragment Rabbit Anti-Rat IgG (H+L) (Jackson Immuno Research Lab, cat#312-0060045) to reduce cross-reactivity with rat IgG. Secondary antibodies used were Immun-Star goat anti-rabbit (GAR)-HRP Conjugate (Cat # 170-5046, or Cat#170-6515; Bio-Rad Laboratories, USA). Clarity Western ECL Substrate (cat #170-5061, Bio-Rad) exposure for 3 min followed secondary exposure. For DA D_1_ quantification, since no standard was available, after development of each blot, the mean image intensity for all samples within each blot was averaged. This average for each blot (roughly 10 blots were run) was normalized against one blot to generate a factor by which to apply to the resulting values from each blot. These values were used in statistical analysis This practice mitigated influence of different image intensities caused by blot processing and imaging.

### Statistics

To analyze locomotor results of FAS and open-field, we used repeated measures 2-way ANOVA and one way ANOVA, respectively, to match baseline performances against subsequent results obtained at the 7 day testing intervals. The FAS results documented changes in forelimb use in paws associated with the lesioned and the intact (unlesioned) side. Locomotor activity in the open-field was analyzed

For the neurochemistry results, time post-operation (7 or 28 days) and operation type (6-OHDA or sham) were used as independent variables. The results obtained from the hemisphere contralateral to the hemisphere of the treatment (sham or 6-OHDA) were paired as within-subjects for each time point in a two-way repeated measures analysis, with between-subject comparisons for time post-operation comparing sides relative to treatment. In the sham group, sources of variation were the treatment (sham-operation), days post-sham-operation, and interaction between these two variables. In the 6-OHDA group, sources of variation were the treatment (6-OHDA lesion), days post-lesion, and interaction between these two variables. Evidence for significant effects in any of these three sources of variation were further analyzed by Student’s t-tests; paired comparisons for within-subjects analysis (in 6-OHDA group, contralateral to lesion vs. lesion within each day post-lesion, and in Sham group, intact vs. sham op), and unpaired comparisons for between-subjects comparisons (in 6-OHDA group, comparison of results from day 7 vs day 28 for each side respective to lesion, and in Sham group, comparison of results from day 7 vs day 28 for each side respective to sham-op). We also determined if the 6-OHDA lesion affected each dependent variable contralateral to lesion by comparing results from the side contralateral to 6-OHDA lesion to the results of the sham-operated side of the sham-operated group.

In the lesioned groups (day 7 and day 28 cohorts), results obtained from rats that did not exhibit greater than 25% loss of forelimb use, as compared to use of the forelimb associated with the non-lesioned hemisphere, were excluded from analyses of neurochemistry and locomotor results. Our results show that more than 25% reduction of forelimb use strongly predicted onset of motor impairment in the open-field. Grubb’s test was applied after excluding values associated with <25% forelimb use to identify outliers, with alpha set to 0.05.

## Results

### Motor function

#### Forelimb use

After nigrostriatal lesion, forelimb use decreased overall to less than 50% versus use on the contralateral side at day 7, in the 7-day cohort (t=7.41, *p* <0.0001, df=8) and 28-day cohort (t=11.77, *p* <0.0001, df=8) (data not shown), with no significant difference between cohorts (Fig. 1A). Decreased forelimb use was maintained 14, 21, and 28 days in the 28-day cohort (Fig. 1B), with no further decrease (F_(3,24)_ =1.80, ns). There was also a significant and sustained increase (15-22% of baseline) in forelimb use associated with the contralateral nigrostriatal pathway (Fig. 1C).

**Figure 1.**
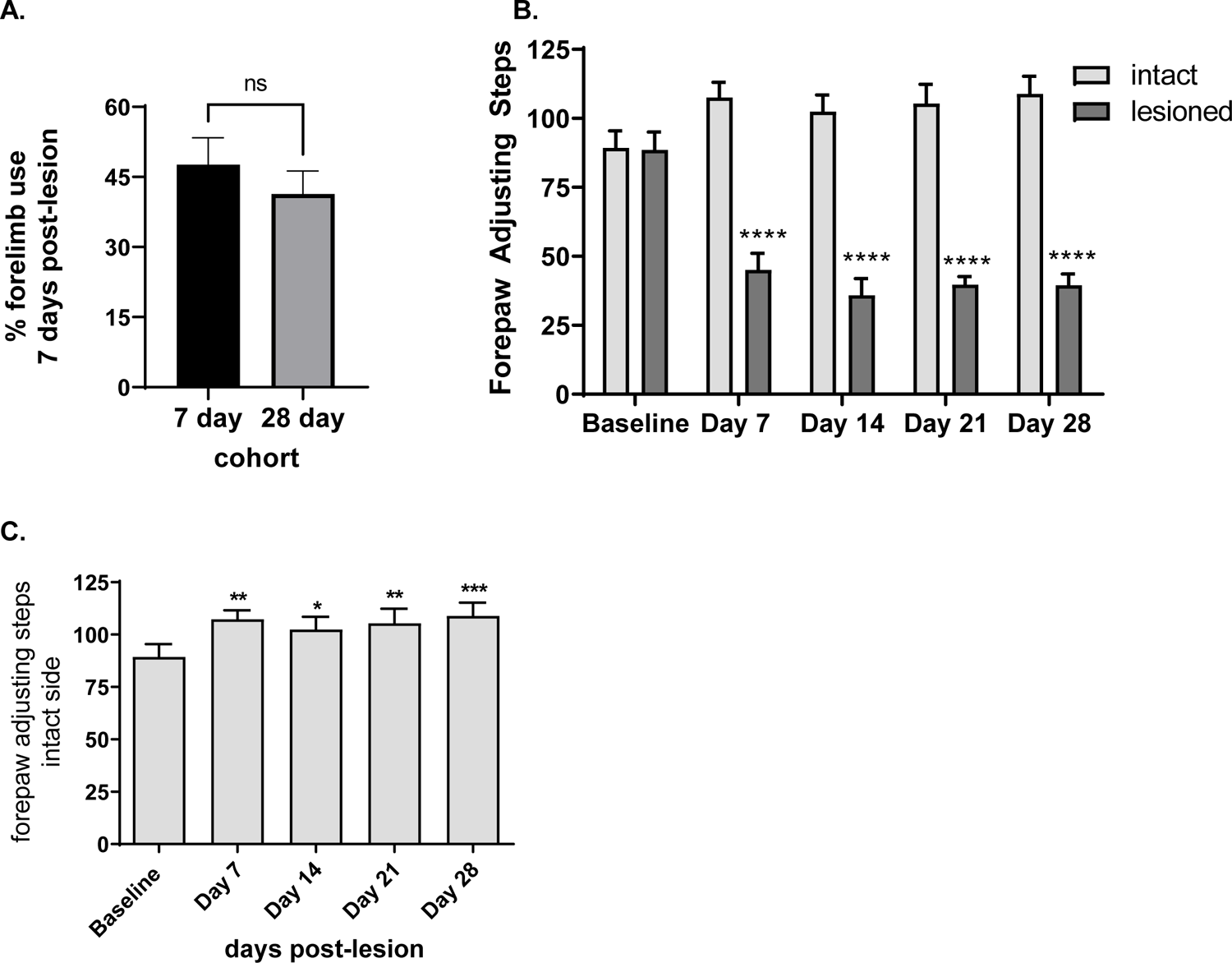
Timeline of forepaw adjustment step differentials of forelimb use associated with lesioned and intact nigrostriatal pathway. **A. Comparison of forelimb use 7 days after nigrostriatal hemi-lesion induction in 7-day vs 28-day cohorts.** No significant difference in use of forelimb associated with nigrostriatal lesion 7 days after lesion between the 7-day and 28-day cohort (t =0.82, ns, df=16). **B. Forelimb use associated with intact vs. lesioned nigrostriatal pathway.** Forelimb use decreased by 7 days following lesion and did not recover for out to 28 days post-lesion (Lesion, F_(1,16)_ = 50.7, *p*<0.0001; time after lesion F_(4,64)_ = 15.5, *p*<0.0001, lesion X time after lesion F_(4,64)_ = 57.8, *p*<0.0001). Baseline (t=0.09, ns); day 7 (t=7.63, \*\*\*\**p<*0.0001); day 14 (t=8.21, \*\*\*\**p<*0.0001); day 21 (t=8.00, \*\*\*\**p<*0.0001); day 28 t=8.45, \*\*\*\**p<*0.0001). **C. Forelimb use associated with intact nigrostriatal pathway.** Forelimb use increased after lesion induction on contralateral side (F_(4,32)_ = 6.06, *p*=0.001). Baseline vs. day 7 (q=4.00, \*\**p*<0.01), day 14 (q=2.91, \**p*<0.05), day 21 (q=3.55, \*\**p*<0.01), day 28 (q=4.35, \*\*\**p*<0.001).

#### Locomotor activity in open-field

Nigrostriatal lesion affected all locomotor parameters as days following 6-OHDA induction increased. In the 7-day cohort, distance travelled decreased overall to near 80% (or a decrease of 20%) of baseline, but the decrease was heterogenous in rats with confirmed >25 % reduction in forelimb use. Just over half of the rats had locomotor activity < 90% of baseline (Fig. 2.A). In the 28-day cohort, locomotor activity uniformly and significantly decreased by 21 days and continued to decline by day 28 (Fig. 2B). There was no significant difference in distance travelled between the 7-day cohort (3697 ± 421) and the 28-day cohort at day 7 (3970 ±371) (data not shown (t=0.49, ns, df=16)), demonstrating comparable lesion induction and progression rate between cohorts.

**Figure 2.**
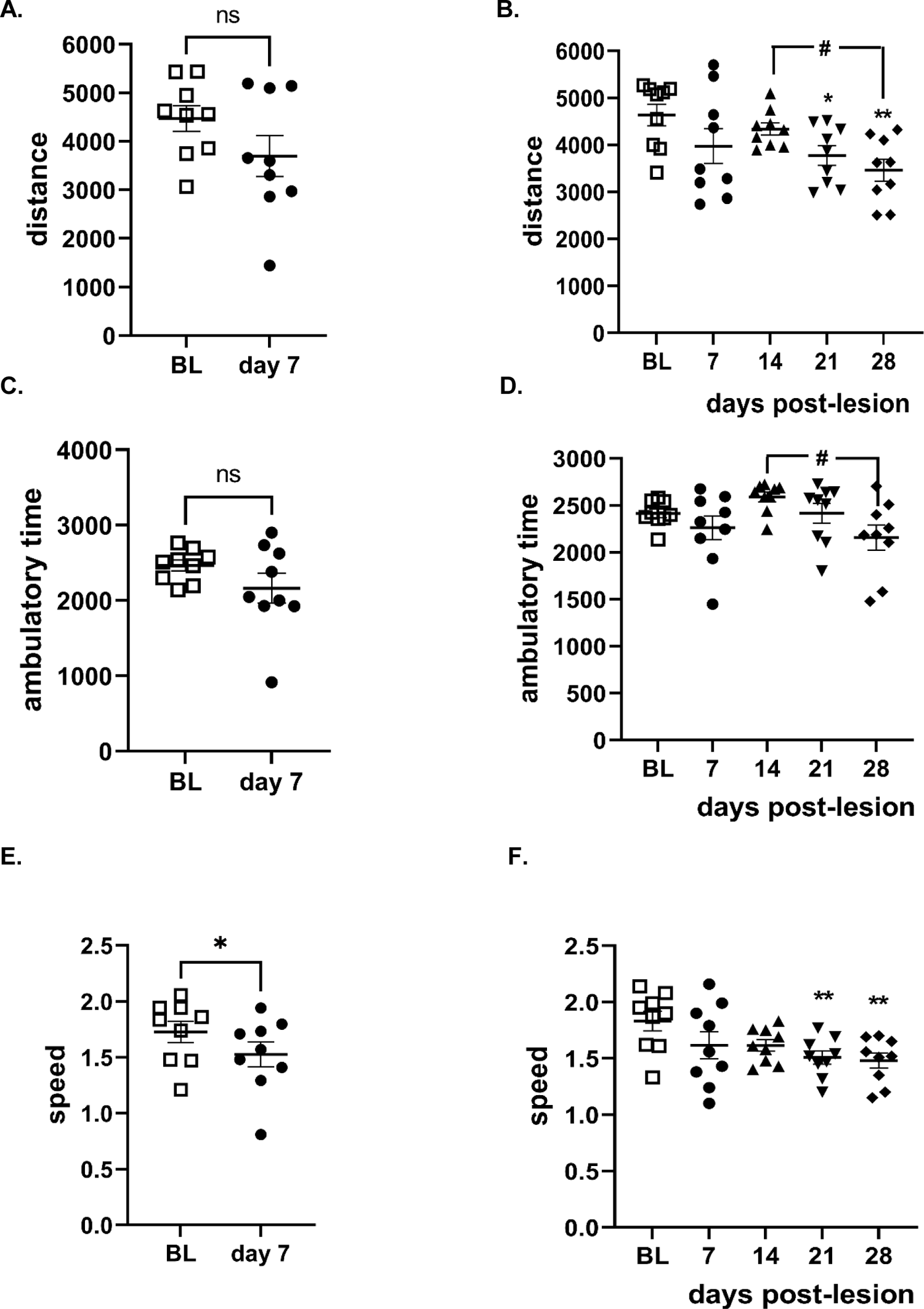
Progressive decline in locomotor activity in open-field following nigrostriatal lesion. **A. Total distance, 7-day cohort.** Total distance was not significantly different between baseline (BL) and day 7 post-lesion (t=2.24, *p*=0.06, df=8). **B. Total distance, 28-day cohort.** Total distance decreased as time post-lesion increased (F_(4,32)_ = 4.94, *p=*0.003); BL v day 7 (q=3.20, ns), BL v day 14 (q=1.44, ns), BL v day 21 (q=4.14, \**p*<0.05), BL v day 28 (q=5.66, \*\**p*=0.003), day 14 v day 28 (q=4.22, ^#^*p*<0.05) Tukey’s multiple comparisons test. **C. Ambulatory time, 7-day cohort.** Time spent moving was not significantly different at 7 days following nigrostriatal lesion (t=1.70, ns, df=8). **D. Ambulatory time, 28-day cohort.** Time spent moving differed as time post-lesion increased (F_(4,32)_ = 3.23, *p=*0.02), but without significant difference between BL and specific post-lesion time. Day 14 v day 28 (q=4.70 ^#^*p*<0.05) Tukey’s multiple comparisons test. **E. Speed, 7-day cohort**. Movement speed decreased ∼12% 7 days following nigrostriatal lesion (t=2.55, \**p*<0.05, df=8). **F. Speed, 28-day cohort.** Movement speed decreased ∼20% (or 80% of baseline speed) as time post-lesion increased. Lesion (F_(4,32)_ = 4.81, *p*=0.004), BL v day 7 (q=3.42, ns), BL v day 14 (q=3.46, ns), BL v day 21 (q=5.14,\*\**p*<0.01), BL v day 28 (q=5.58, \*\**p*<0.001), Tukey’s multiple comparisons test.

Time spent moving (ambulatory time) decreased over time following 6-OHDA lesion induction with confirmed reduced forelimb use at day 7, with no significant decrease 7 days after lesion induction (Fig. 2C), but a significant decrease by day 28 post lesion (Fig. 2D). There was no significant difference in this parameter at day 7 between the 7-day cohort (2162 ± 199) and the 28-day cohort at day 7 (2260 ± 128) (data not shown (t=0.42, ns, df=16).

Movement speed was slightly (∼12%) but significantly decreased in the 7-day cohort (Fig. 2E), and in the 28-day cohort, there was a progressive decline in speed, reaching significance by day 21 (Fig. 2F). There was no significant difference in speed between the 7-day cohort (1.53 ± 0.11) and the 28-day cohort at day 7 (1.61 ± 0.12) (data not shown (t=0.55, ns, df=16)), indicating comparable lesion induction between cohorts, also observed in the other open-field parameters.

The severity of forelimb use deficiency predicted eventual 6-OHDA-induced decreases in locomotor activity. Rats with <25% reduction in FAS 7 days after lesion did not exhibit decreased locomotor activity at any time post-lesion, whereas rats with >25% FAS reduction exhibited decreased locomotor activity between day 21 to 28 (Fig. 3).

**Figure 3.**
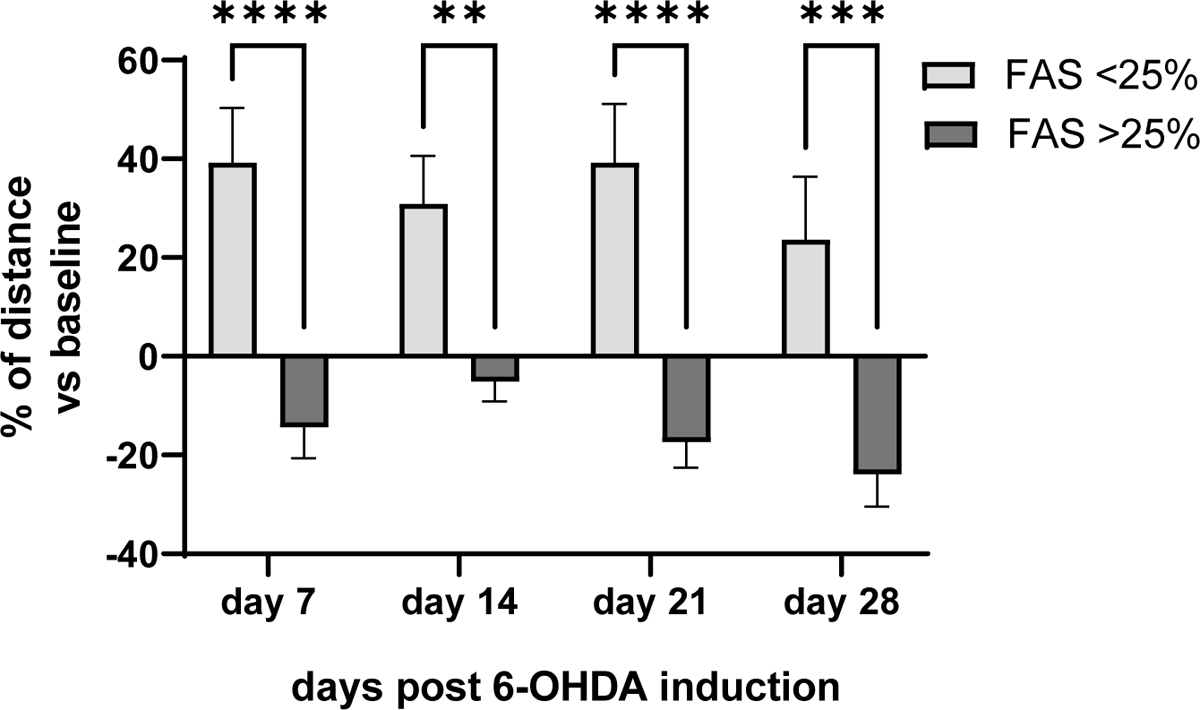
Critical threshold of reduced forelimb use for progressive decline in locomotor activity. Forelimb use associated with lesioned nigrostriatal pathway at 7 days post-lesion predicts eventual decrease in locomotor activity. Rats were evaluated for FAS on day 7 post-6-OHDA delivery into the medial forebrain bundle and assigned into two groups; FAS less than 25% reduction (*n=*4) and FAS greater than 25% reduction of forelimb use (*n*=9). Forelimb impairment severity at the 25% reduction threshold was significantly associated with eventual decrease in locomotor activity (F_(1,11)_=31.3, *p*=0.0002). Day 7 (t=4.78, \*\*\*\**p*<0.0001); Day 14 (t=3.21, \*\**p*<0.0001); Day 21 (t=5.05, \*\*\*\**p*<0.0001); Day 28 (t=4.24, \*\*\**p=*0.0005).

### Tyrosine hydroxylase vs total neuron loss

6-OHDA lesion progressively decreased TH+ neuron number (Fig. 4A,C). With loss of TH protein in striatum confirmed to be ≥ 80% (By cohort; 7-day, 93 ± 4%; 28-day, 98 ± 1%), there was a highly significant effect of lesion, days post lesion, and interaction. Notably, 7 days following lesion, TH cell loss varied considerably, ranging from 12 to 44% vs. TH cell number contralateral to lesion (Fig. 4A), and this variability was also seen in the total neuron loss (Nissl stain) ranging from 7 to 21% (Fig. 4B). However, by day 28 after lesion induction, variability was reduced nearly 4-fold, with TH cell loss ranging from 70-94% and total neuron loss ranging from 39-62% (Fig. 4A, B). This variability in TH cell loss was restricted to the lesioned side, as TH cell numbers were 2- to 3-fold less variable contralateral to lesion at both time points.

**Figure 4.**
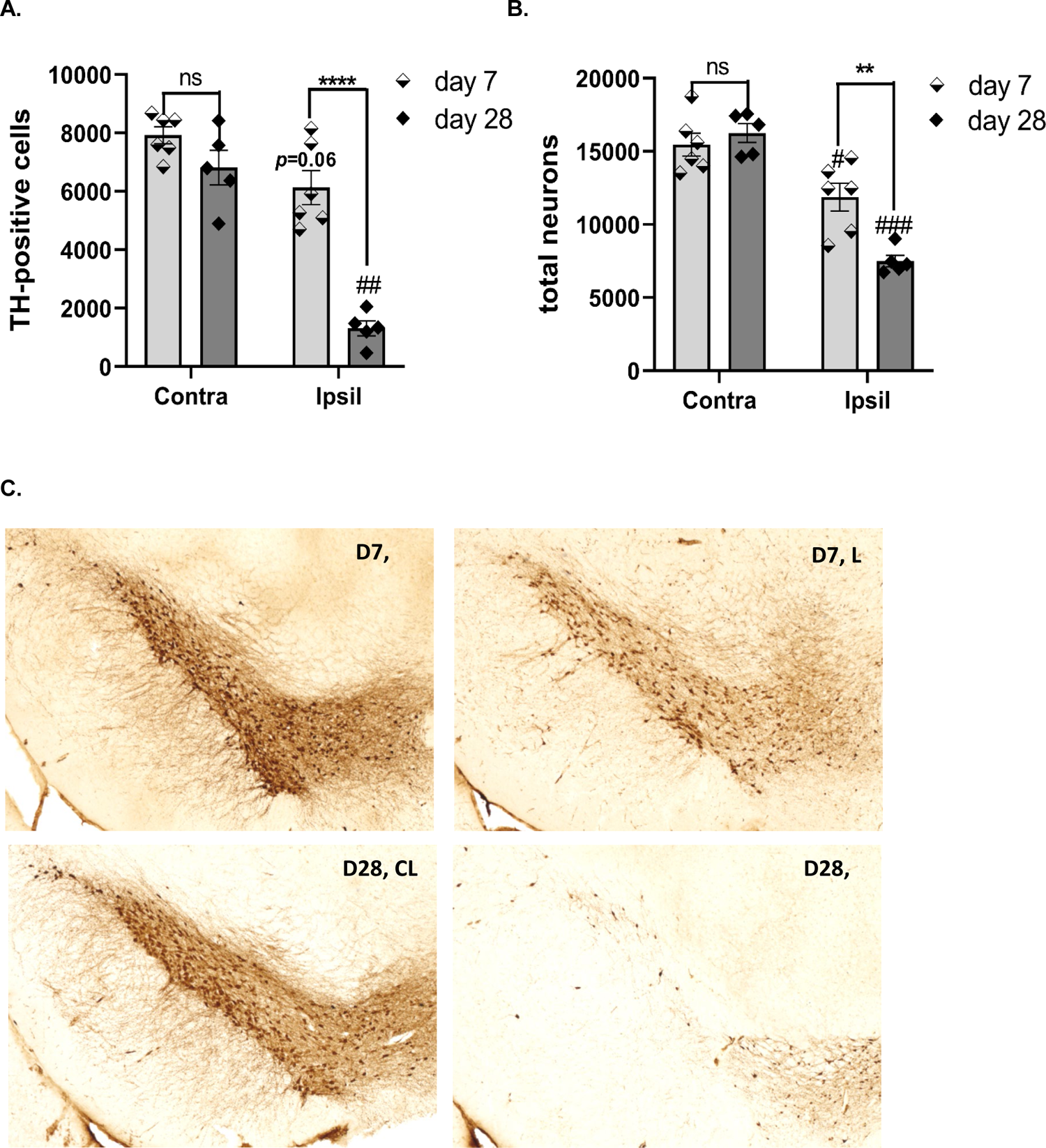
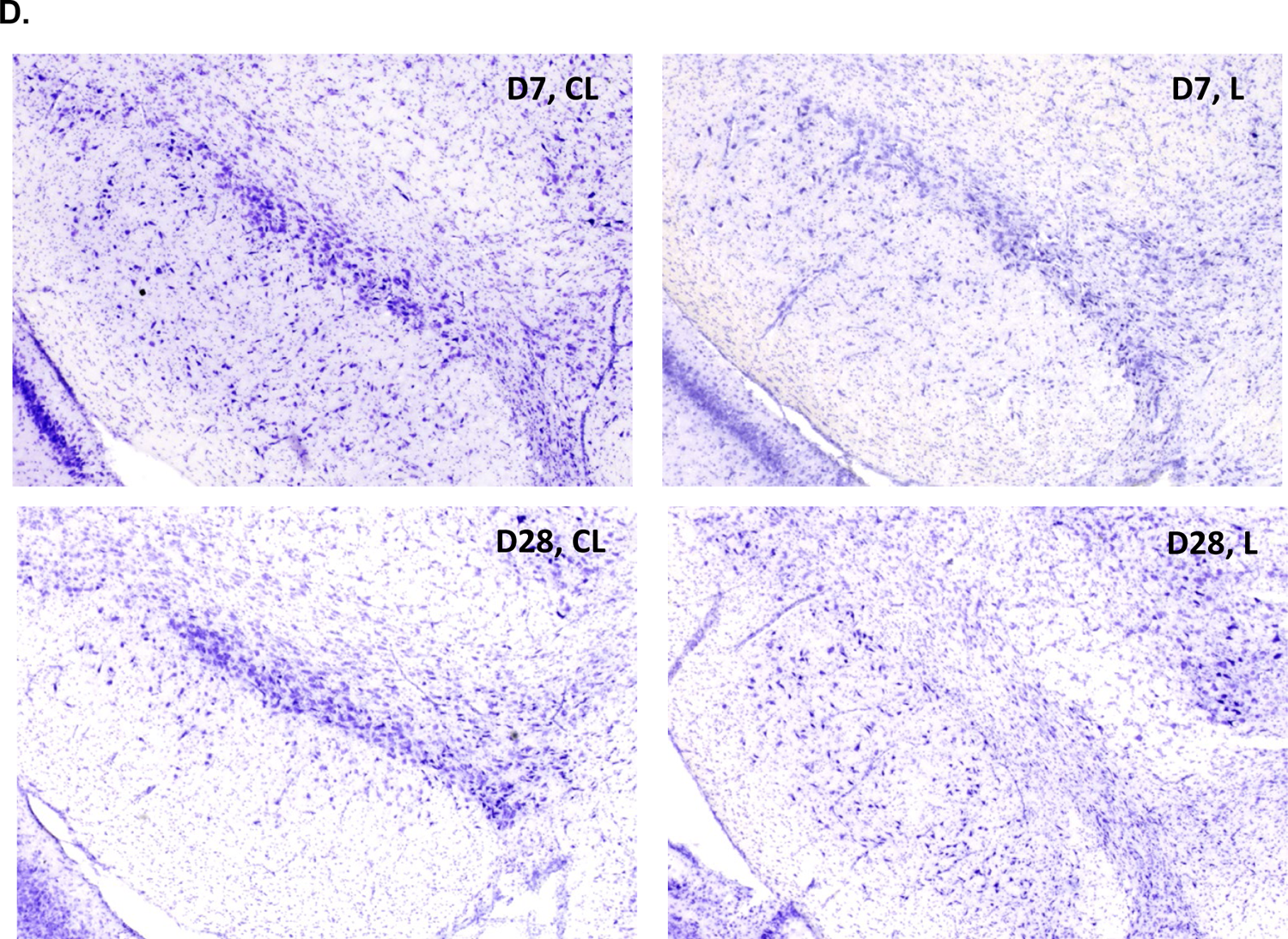
Timeline of nigrostriatal neuron and cell loss following 6-OHDA lesion induction. Results reflect a minimum of 25% reduction in forelimb use. **A. TH+ neuron loss.** TH+ neuron loss occurred after lesion induction and was progressive between day 7 and 28. Lesion (F_(1,9)_ = 51.4, *p*<0.0001); days post-lesion (F_(1,9)_ = 50.8, *p*<0.0001); lesion x days post-lesion (F_(1,9)_ = 13.3, *p=*0.005). Day 7 vs day 28; lesioned side (t=7.02, \*\*\*\**p*<0.0001, df=9); contralateral to lesioned side (t=1.77, ns, df=9). Contra vs Ipsil to lesion; Day 7 (t=2.43, *p=*0.06, df=5); Day 28 (t=8.16 ^##^*p=*0.001,df=4). **B. Total neuron loss (Nissl).** Total neuron loss followed lesion induction and was progressive between day 7 and 28. Lesion (F_(1,9)_ = 74.3, *p*<0.0001); days post-lesion (F_(1,9)_ = 4.93, *p*=0.054); lesion x days post-lesion (F_(1,9)_ = 13.0, *p=*0.006). Day 7 vs day 28; lesioned side (t=3.92, \*\**p*=0.0035, df=9); contralateral to lesioned side (t=0.75, ns, df=9). Contra vs Ipsil to lesion; Day 7 (t=3.39, ^#^*p=*0.02, df=5); Day 28 (t=9.59 ^###^*p=*0.0007, df=4). **C. Representative image of TH cell loss over time.** Left panels, contralateral to lesion (CL); right panels; ipsilateral to lesion (L). Top panels, Day 7; bottom panels, Day 28. **D. Representative image of total cell loss cell loss over time** Left panels, contralateral to lesion; right panels; ipsilateral to lesion. Top panels, Day 7; bottom panels, Day 28

Thus, TH cell loss was progressive between 7- and 28-days post-lesion, with possible resilience evident in some rats against loss early (day 7) after lesion induction, despite confirmed >90% TH loss in striatum. TH cell loss exceeded total neuron loss by day 28 after lesion induction, which may be attributable to a decrease in TH neuron number or TH protein, or combination.

### Dopamine and dopamine turnover

The rate of DA loss between the striatum and SN progressed at different rates after lesion induction. By day 7, there was maximal DA loss (>95%) in striatum, sustained out to day 28 (Fig. 5A). However, in the SN, DA loss was more protracted, with no overall loss at day 7, but significant loss (64-72%) by day 28 (Fig. 5B).

**Figure 5.**
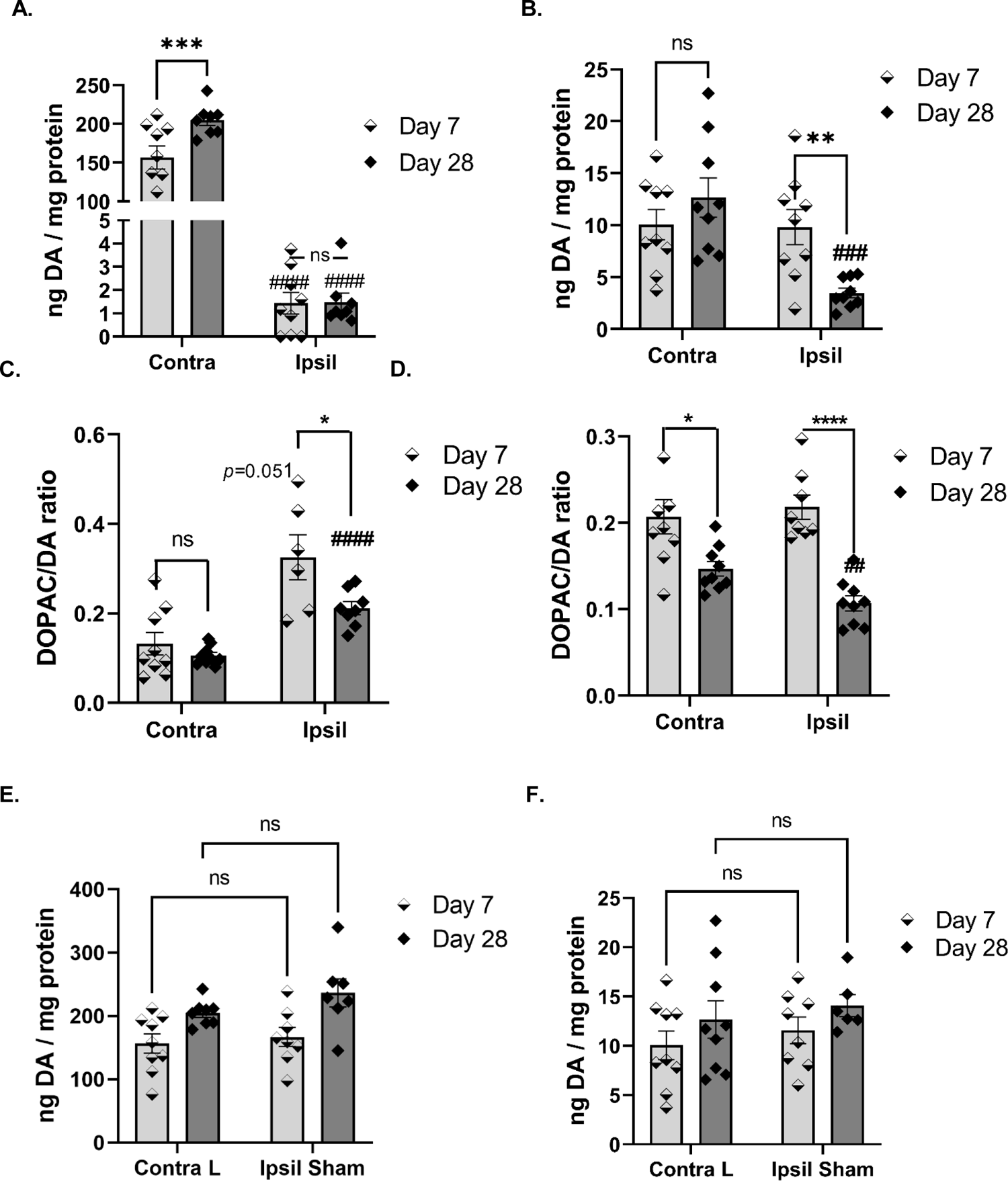
Disparity in rate of nigrostriatal DA loss and turnover in nigrostriatal pathway. **A. DA loss, Striatum.** DA loss was maximal on or before 7 days with FAS-confirmed lesion. Lesion (F_(1,15)_ = 418, *p*<0.0001); days post-lesion (F_(1,15)_ = 7.93, *p*=0.013); lesion x days post-lesion (F_(1,15)_ = 7.51, *p=*0.015). Day 7 vs day 28; lesioned side (t=0.085, ns, df=15); contralateral to lesioned side (t=3.12, \*\**p* <0.01, df=16). Contra vs Ipsil to lesion; Day 7 (t=10.1, ^####^*p* <0.0001,df=8); Day 28 (t=28.3, ^####^*p* <0.0001,df=7). **B. DA loss, SN.** Lesion (F_(1,16)_ = 12.9, *p*=0.0025); days post-lesion (F_(1,16)_ = 1.31, ns); lesion x days post-lesion (F_(1,16)_ = 11.71, *p=*0.005). Day 7 vs day 28; lesioned side (t=3.62, \*\**p* <0.01, df=16); contralateral to lesioned side (t=1.10, ns, df=16). Contra vs Ipsil to lesion; Day 7 (t=0.12, ns, df=8); Day 28 (t=5.2, ^###^*p* <0.001, df=8). **C. DA turnover, Striatum.** DA loss was maximal on or before 7 days with FAS-confirmed lesion. Lesion (F_(1,28)_ = 35.7, *p*<0.0001); days post-lesion (F_(1,28)_ = 7.86, *p*=0.009); lesion x days post-lesion (F_(1,28)_ = 3.04, ns). Day 7 vs day 28; lesioned side (t=2.48, \**p*<0.05, df=12); contralateral to lesioned side (t=1.01, ns, df=16). Contra vs Ipsil to lesion; Day 7 (t=2.56, *p*=0.051, df=5); Day 28 (t=8.0, ^####^*p* <0.0001, df=7). **D. DA turnover SN.** Lesion (F_(1,31)_ = 1.09, ns); days post-lesion (F_(1,31)_ = 39.6, ^####^*p* <0.0001); lesion x days post-lesion (F_(1,31)_ = 3.52, *p=*0.07). Day 7 vs day 28; lesioned side (t=6.86, \*\*\*\**p* <0.0001, df=15); contralateral to lesioned side (t=2.79, \**p*<0.05, df=16). Contra vs Ipsil to lesion; Day 7 (t=0.71, ns, df=7); Day 28 (t=4.13, ^##^*p* <0.01, df=8). **E. Str, contra to lesion vs sham-op.** 6-OHDA hemi-lesion did not affect DA levels in striatum contralateral to lesion at either day post-lesion. Contra to lesion vs sham-op (F_(1,28)_ = 1.87, ns); sham-operation x days post sham-operation (F_(1,28)_ = 0.49, ns). There was a difference in levels of DA at day 28 v day 7 in both contra to lesion in the 6-OHDA group and the sham-op (F_(1,28)_ = 14.76, *p* <0.001). Based on results from the sham-operation (Suppl Fig S6A), this increase was likely due to the effect of sham-operation over time. **F. SN, contra to lesion vs sham-op.** 6-OHDA hemi-lesion did not affect DA levels in SN contralateral to lesion at either day post-lesion. Contra to lesion vs sham-op (F_(1,28)_ = 0.86, ns); sham-operation x days post sham-operation (F_(1,28)_ = 0.001, ns), or time post-lesion (F_(1,28)_ = 2.65, ns).

Nigrostriatal lesion affected DA turnover differently between the striatum and SN. In striatum, in conjunction with >95% DA loss, DA turnover increased in the lesioned side at both day 7 and day 28 (Fig. 5C). Notably, the increase in DA turnover by day 28 was significantly less versus day 7, although still elevated compared to contralateral striatum (Fig. 5C). In the SN, the time past lesion induction had a highly significant effect on DA turnover in both lesioned and contralateral to lesion sides (Fig. 5D).

In the mesoaccumbens pathway, there was ∼90% loss of DA in the NAc, with maximal loss by day 7 after lesion induction, similar to striatum (Suppl Fig. 6A). However, in VTA, DA increased by day 7, and then receded to levels observed in contralateral to lesion at both time points by day 28 (Suppl Fig. 6B).

Sham-operation produced a time-dependent difference in DA tissue levels between day 7 and day 28 in striatum (Suppl Fig. 7A), but had no effect in SN (Suppl Fig. 7B). In the mesoaccumbens pathway, there was a time-related difference in DA levels in the NAc (Suppl Fig. 7C), but no effect in VTA (Suppl Fig. 7D).

### Dopamine loss vs. open-field activity

7 days after lesion induction in either cohort, there was considerable heterogeneity in the change of locomotor activity from baseline by day 7, ranging from −53% to +14% in the day 7 cohort (mean +/- SD, −18.0 +/- 23.8 %) and −44% to +8% in the day 28 cohort (mean +/- SD, −14.4 +/- 18.6%). This variability from baseline was still present by day 28, ranging from −51 to + 20% relative to baseline (mean +/- SD = −24 +/- 20 %). Thus, despite confirming reduced forelimb use by day 7 (Fig. 1A,B, 3), the impact of 6-OHDA lesion on locomotor activity was variable, suggesting plasticity in DA signaling in response to nigrostriatal neuron loss was also variable. We evaluated if differences in striatal or nigral DA loss at day 7 and 28 aligned against the change in baseline. Striatal DA loss had no correlation with the change in baseline activity (Fig. 6A). However, the magnitude of nigral DA loss was highly correlative with the magnitude of decrease from baseline locomotor activity (Fig. 6B).

**Figure 6.**
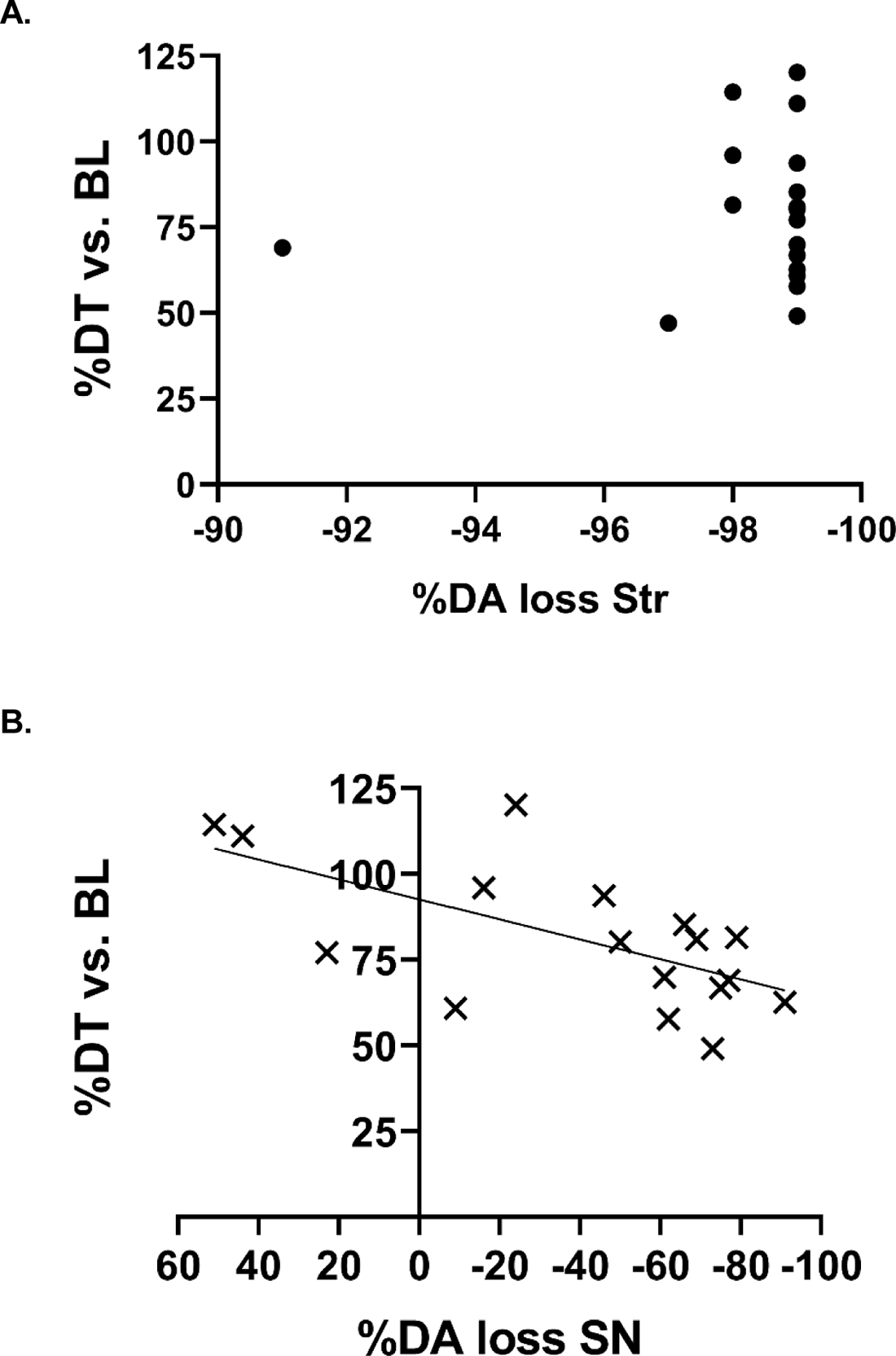
Relationship between striatal and nigral DA loss against motor function parameters. DA loss in the SN, but not striatum, corresponded with the % decrease in locomotor activity (as distance traveled (DT)) at day 7 and day 28 against pre-lesion baseline levels. All results are from rats that exceeded >25% reduction in forelimb use by day 7 post-lesion. **A. DA loss in striatum vs. change in locomotor activity.** No correlation between striatal DA loss against the % decrease in locomotor activity. Pearson r = −0.130, R^2^= 0.017, *p* =0.61, 18 XY pairs. **B. DA loss in SN vs. change in locomotor activity.** There was a significant correlation between nigral DA loss against the % decrease in locomotor activity. Pearson r = 0.629, R^2^= 0.395, *p* =0.007, 17 XY pairs.

### Tyrosine hydroxylase protein expression

TH protein loss in striatum proceeded at a greater rate versus SN after lesion. In the striatum, TH protein loss 7 days post-lesion exceeded 95%, with only ∼4% of TH protein (∼0.02 ng TH/ ug protein) remaining vs levels contralateral to lesion (∼0.40 ng TH/ug protein) (Fig. 7A), and no further decrease by day 28. There was no difference in TH expression contralateral to lesion (Fig. 7A), nor against levels in the sham-operated striatum in the Sham group (Fig. 7C).

**Figure 7.**
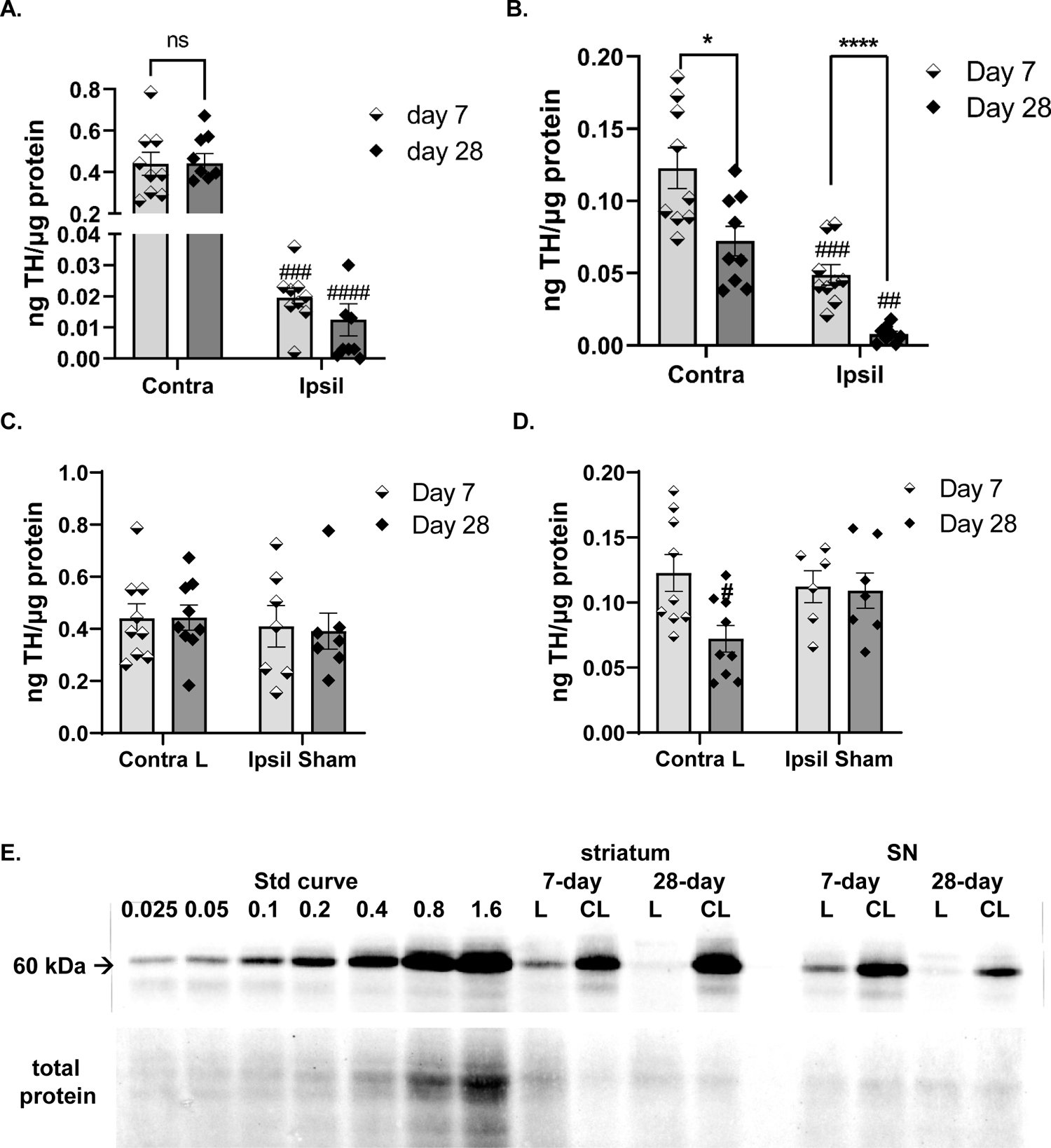
Disparity in rate of TH protein loss in nigrostriatal pathway in lesioned and contralateral to lesioned sides. A-C, 6-OHDA group only; D, E, Contra to lesion vs sham-op. **A. Striatum.** TH loss was maximal on or before 7 days with FAS-confirmed lesion. Lesion (F_(1,16)_ = 129.6, *p*<0.0001); days post-lesion (F_(1,16)_ = 0.004, ns); lesion x days post-lesion (F_(1,16)_ = 0.016, ns). Day 7 vs day 28; lesioned side (t=0.62, ns, df=15); contralateral to lesioned side (t=0.10, ns, df=16). Contra vs Ipsil to lesion; Day 7 (t=7.46, ^###^*p* =0.0001, df=7); Day 28 (t=8.59, ^####^*p* <0.0001, df=8). **B. SN.** TH protein loss was progressive in SN in lesioned and contralateral to lesion sides. Lesion (F_(1,15)_ = 62.4, *p<*0.0001); days post-lesion (F_(1,16)_ = 17.9, *p=*0.0006); lesion x days post-lesion (F_(1,15)_ = 0.35, ns). Day 7 vs day 28; lesioned side (t=5.24, \*\*\*\**p* <0.0001, df=15); contralateral to lesioned side (t=2.90, \**p*<0.05, df=16). Contra vs Ipsil to lesion; Day 7 (t=5.65, ^###^*p* =0.0005, df=8); Day 28 (t=5.34, ^##^*p* <0.01, df=7). **C. Striatum, contra to lesion.** TH protein loss was unaffected in striatum in side contralateral to lesioned side. Contra to Lesion (F_(1,28)_ = 0.44, ns); days post-lesion (F_(1,28)_ = 0.90, ns); contra lesion x days post-lesion (F_(1,28)_ = 0.03, ns). **D. SN, contra to lesion.** TH protein loss was progressive in SN in side contralateral to lesioned side as a function of time past lesion. Lesion (F_(1,27)_ = 1.03, ns); days post-lesion (F_(1,27)_ = 4.27, *p=*0.049); contra lesion x days post-lesion (F_(1,27)_ = 3.36, *p* =0.08). Day 7; contralateral to lesioned side vs Sham (t=0.53, ns, df=13). Day 28; contralateral to lesioned side vs Sham (t=2.22, \**p<*0.05, df=14). **E. Representative quantitative blot images of TH protein expression in striatum and SN.** Total protein of 5 or 10 µg was loaded to quantify TH protein in lesioned or contralateral to lesioned striatum and SN. The standard curve ranged from 0.025 to 1.6 ng total TH protein. TH protein expression in the lesioned (L) or contralateral to lesioned (CL) striatum and SN are represented from the 7-day cohort and the 28-day cohort. 10 µg of total protein was loaded from lesioned striatum samples, and 5 µg total protein was loaded from the contralateral to lesioned striatum samples. 5 µg total protein was loaded from all SN samples.

In contrast, TH protein loss in the SN was more protracted, with 36% remaining [(0.05 ng TH/ug protein) vs contralateral SN (0.12 ng TH/ug protein)] on day 7, progressing to 11% TH remaining [(∼0.01 ng TH/ ug protein) vs contralateral SN (∼0.07 ng TH/ ug protein)] by day 28 (Fig. 7B). Moreover, there was time-dependent and significant decrease in TH protein expression in the SN contralateral to the lesioned hemisphere, with no loss at day 7, but reduced by ∼40% by day 28 (Fig. 7B); a difference that held when also compared against TH levels in the Sham group (Fig. 7D).

There was no correlation between TH protein loss between the striatum and SN at 7 days (r=0.195, *p* =0.62) or 28 days (r=0.219, *p* =0.22) post-lesion induction (data not shown), indicating that processes governing TH protein loss differed between the stratum and SN.

In the NAc and VTA, TH loss in the NAc proceeded at a much faster rate (Suppl Fig 9A) than in the VTA (Suppl Fig 9B) and was progressive. In the VTA, there was no loss of TH by day 7, but significant loss in the lesioned side by day 28 (Suppl Fig 9B). Thus, TH loss in VTA proceeded at a slower rate than in SN at equivalent time points. Sham-operation did not affect TH protein levels at either time point in either striatum or SN (Suppl Fig 8 A,B).

### TH phosphorylation

Comparison of remaining DA vs TH protein following lesion (as percentage of side contralateral to lesion) revealed that DA remaining was 3-fold less than TH remaining in the striatum (1.5% vs 5% (Fig. 8A). The opposite was observed in the SN, with 2-fold greater DA than TH remaining at both time points after lesion (Fig. 8B), suggesting increased TH activity could be offset by increased TH phosphorylation stoichiometry (PS) in the SN. Although ser40 TH is reportedly found to increase in striatal regions in post-mortem tissue of PD patients [35], *in vivo* evidence indicates no correlation between ser40 TH phosphorylation stoichiometry vs inherent DA tissue levels [31,45–47]. TH phosphorylation stoichiometry at ser40 was unchanged in striatum or the SN following lesion at any time (Fig. 8 C-E), or by sham-operation (Suppl. Fig.10 A,D).

**Figure 8.**
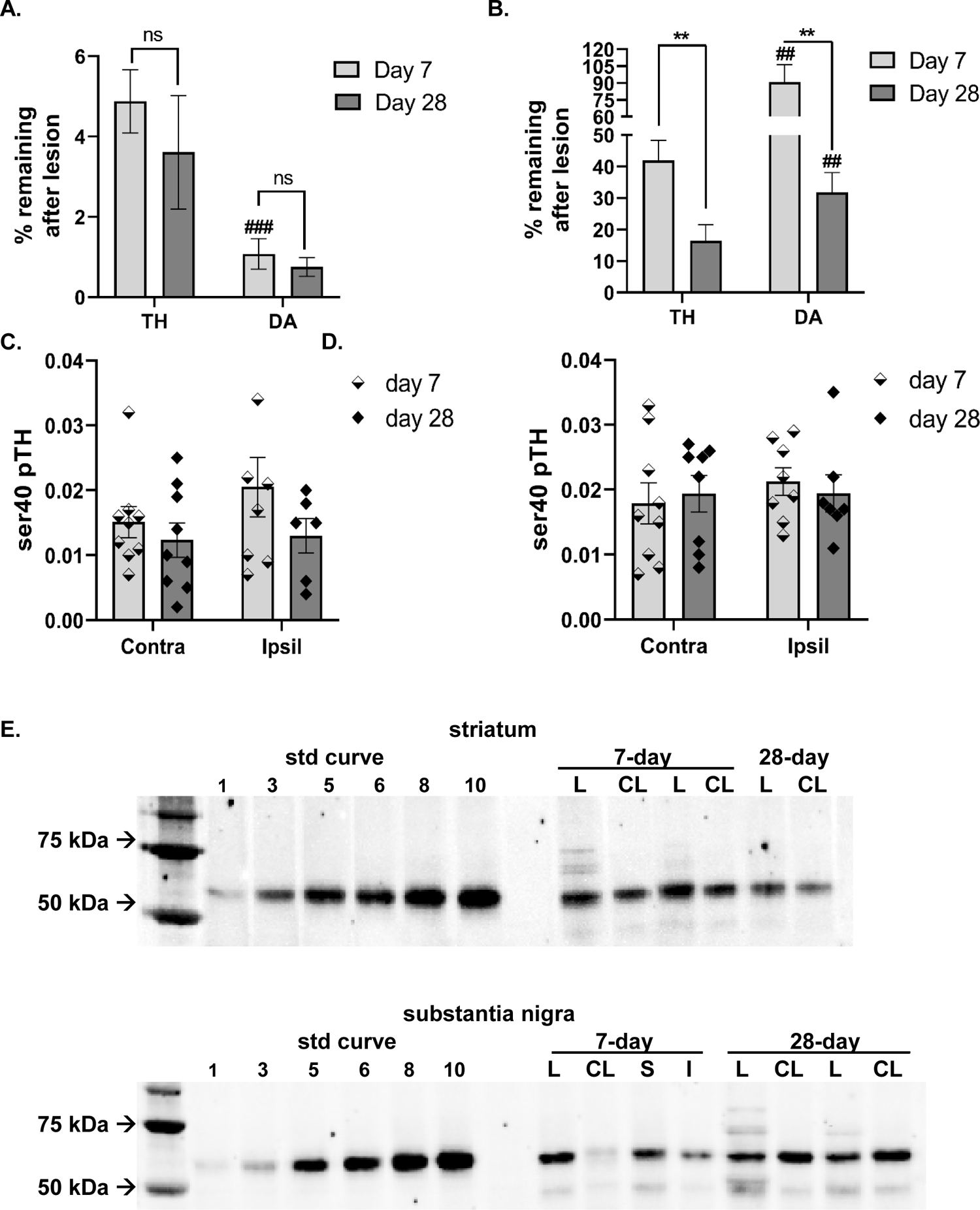
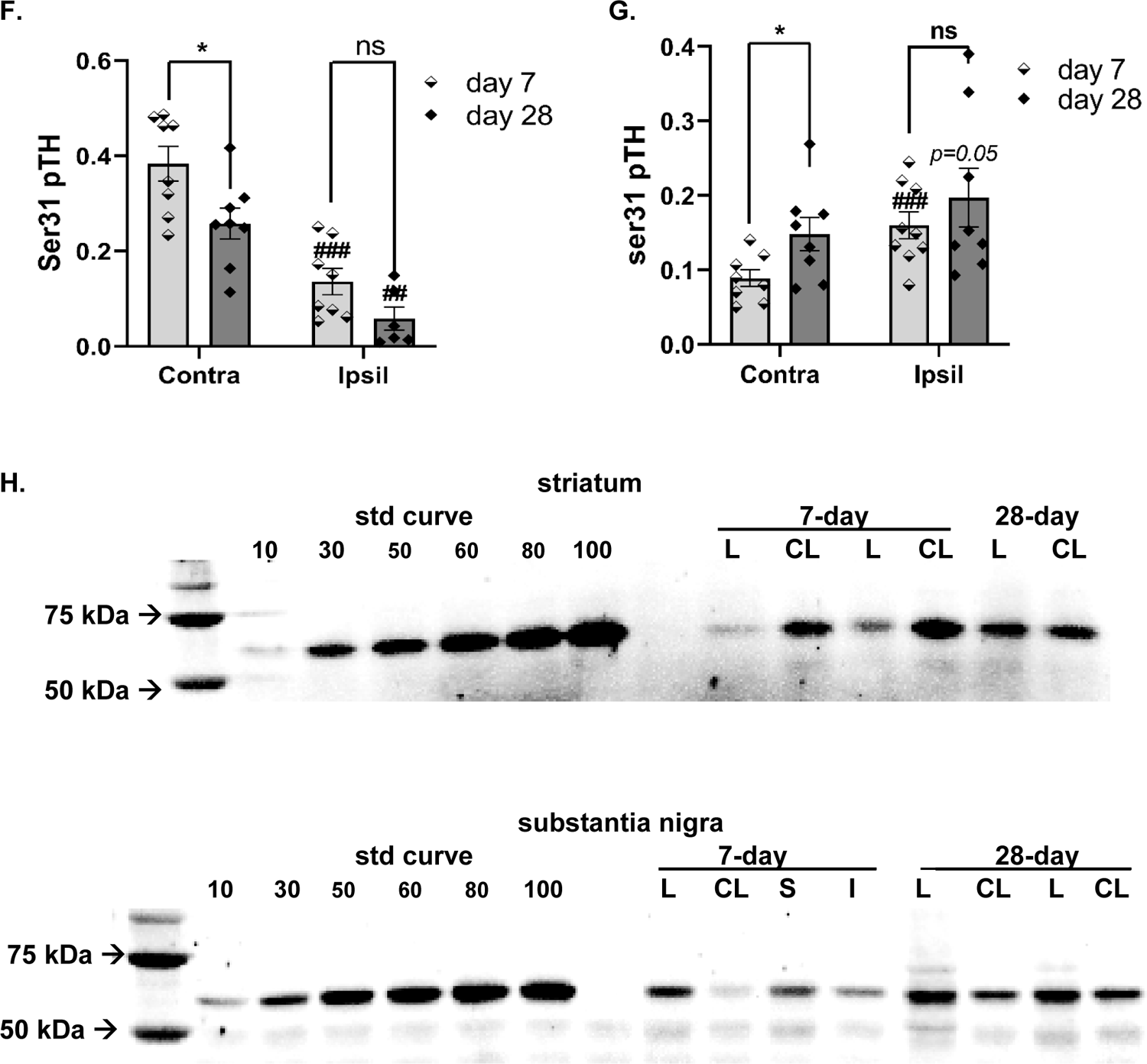
DA loss vs TH protein loss and TH phosphorylation in nigrostriatal pathway in lesioned and contralateral to lesioned sides. A,B DA vs TH protein loss; C-E, Ser31 TH phosphorylation; F-H, Ser40 TH phosphorylation. **A. Striatum.** There was a highly significant difference in DA vs TH remaining after lesion induction (F_(1,15)_ = 18.4, p=0.0007), but no effect of time after lesion between 7 or 28 days after lesion; days post-lesion (F_(1,16)_ = 0.47, ns); or interaction, DA v TH remaining x days post-lesion (F_(1,15)_ = 0.63, ns). Day 7 vs day 28; TH (t=0.78, ns, df=16), DA (t=0.72, ns, df=15). DA vs TH remaining; Day 7 (t=5.26, ^###^*p* =0.0008, df=8); Day 28 (t=1.64, ns, df=7). **B. SN.** There was a highly significant difference in DA vs TH remaining after lesion induction (F_(1,15)_ = 27.7, *p* <0.0001), and effect of time after lesion between 7 and 28 days after lesion; days post-lesion (F_(1,16)_ = 15.84, *p* =0.001); interaction, DA v TH remaining x days post-lesion (F_(1,15)_ = 7.54, *p* = 0.015). Day 7 vs day 28; TH (t=3.12, *p*=0.007, df=16), DA (t=3.76, *p=*0.002, df=15). DA vs TH remaining; Day 7 (t=3.99, ^##^*p* =0.005, df=7); Day 28 (t=3.42, ^##^*p* =0.009, df=7). **C. Ser40 Striatum.** Ser40 TH phosphorylation in striatum was unaffected by nigrostriatal lesion. Lesion (F_(1,12)_ = 1.21, ns); days post-lesion (F_(1,16)_ = 1.99, ns); lesion x days post-lesion (F_(1,12)_ = 1.33, ns). D. **Ser40 SN.** Ser40 TH phosphorylation in SN was unaffected by nigrostriatal lesion. Lesion (F_(1,12)_ = 0.99, ns); days post-lesion (F_(1,16)_ = 0.01, ns); lesion x days post-lesion (F_(1,12)_ = 0.50, ns). Striatum. E. Representative quantitative blot images of Ser40 TH PS in striatum and SN. Top, striatum. 0.2 ng TH protein was loaded for each sample against a standard curve of 1 to 10 pg of phosphorylated ser40 standard. Lesioned (L), Contralateral to lesioned (CL) are represented in two rats from the 7-day cohort and one rat from the 28-day cohort. **Bottom, SN.** 0.2 ng TH protein was loaded for each sample against a standard curve of 1 to 10 pg of phosphorylated ser40 standard. Lesioned (L), Contralateral to lesioned (CL) and sham-op (S) and associated intact side (I) are represented in one rat in the 6-OHDA and sham-operated group from the 7-day cohort and two rats from 6-OHDA 28-day cohort. F. **Ser31 Striatum.** Ser31 TH phosphorylation decreased in striatum following nigrostriatal lesion. Lesion (F_(1,12)_ = 99.3, *p*<0.0001); days post-lesion (F_(1,15)_ = 7.49, *p* =0.015); lesion x days post-lesion (F_(1,11)_ = 0.71, ns). Day 7 vs day 28; lesioned side (t=2.01, *p*=0.07, df=12); contralateral to lesioned side (t=2.59, \**p*=0.02, df=14). Contra vs Ipsil to lesion; Day 7 (t=7.40, ^###^*p* =0.0003, df=6); Day 28 (t=5.80, ^##^*p* =0.002, df=5). **G. Ser 31 SN.** Ser31 TH phosphorylation increased in SN following nigrostriatal lesion. Lesion (F_(1,14)_ = 22.8, *p=*0.0003); days post-lesion (F_(1,15)_ = 2.39, ns); lesion x days post-lesion (F_(1,14)_ = 0.91, ns). Day 7 vs day 28; lesioned side (t=0.89, ns, df=15); contralateral to lesioned side (t=2.35, \**p*=0.03, df=14). Contra vs Ipsil to lesion; Day 7 (t=4.97, ^##^*p* =0.002, df=7); Day 28 (t=2.32, *p* =0.05, df=7). **H. Representative quantitative blot images of Ser31 TH PS in striatum and SN. Top, striatum.** 0.1 ng TH protein was loaded for each sample against a standard curve of 10 to 100 pg of phosphorylated ser31 standard. Lesioned (L), Contralateral to lesioned (CL) are represented in two rats from the 7-day cohort and one rat from the 28-day cohort. **Bottom, SN.** 0.2 ng TH protein was loaded for each sample against a standard curve of 10 to 100 pg of phosphorylated ser31 standard. Lesioned (L), Contralateral to lesioned (CL) and sham-op (S) and associated intact side (I) are represented in one rat in the 6-OHDA and sham-operated group from the 7-day cohort and two rats from 6-OHDA 28-day cohort.

The sham-operation had minimal, if any, effect on ser31 (Suppl. Fig 10). 6-OHDA lesion produced opposing changes in ser31 phosphorylation between striatum and SN. There was a progressive decrease in striatum ipsilateral to lesion at 7 and 28 days after lesion (Fig. 8F), and also a decrease contralateral to lesion between day 7 and 28 (Fig. 8 F,H). In contrast, ser31 TH PS increased in the lesioned SN (Fig. 8G,H), with more than 50% increase (contralateral ∼0.090) vs ipsilateral to lesion (∼0.160) by day 7 and up to 0.197 ser31 PS by day 28 (Fig. 8G). Notably, ser31 PS also increased >50% contralateral to lesion between day 7 and 28 (Fig. 8G). As DA loss was less than TH loss at day 7 and 28 in lesioned SN, and in the contralateral SN on day 28, these results indicate increased ser31 TH phosphorylation in SN mitigates DA loss against TH loss.

### Norepinephrine

As, the SN receives significant noradrenergic innervation from the locus coeruleus and other nuclei [48, 49], increased NE levels after 6-OHDA could suggest that changes in TH phosphorylation in SN could originate from NE terminals in SN. However, 6-OHDA lesion did not affect NE tissue content in the SN (Suppl Fig 11). Thus, increased ser31 TH phosphorylation in the SN was likely unrelated to any change within noradrenergic terminals.

### DA D_1_ receptor expression

Released DA activates D_1_ receptors on striatonigral neurons to promote GABA release [39, 50], which disinhibits of basal ganglia output to promote movement initiation [37]. In striatum, lesion affected expression of the 75 kDa form, although no significant differences were observed in any post-hoc comparison (Fig. 9A). No differences were observed for the 75kDa form in the SN (Fig. 9B). Sham-operation affected expression of the 75 kDa form in both regions (Suppl. Fig 12).

**Figure 9.**
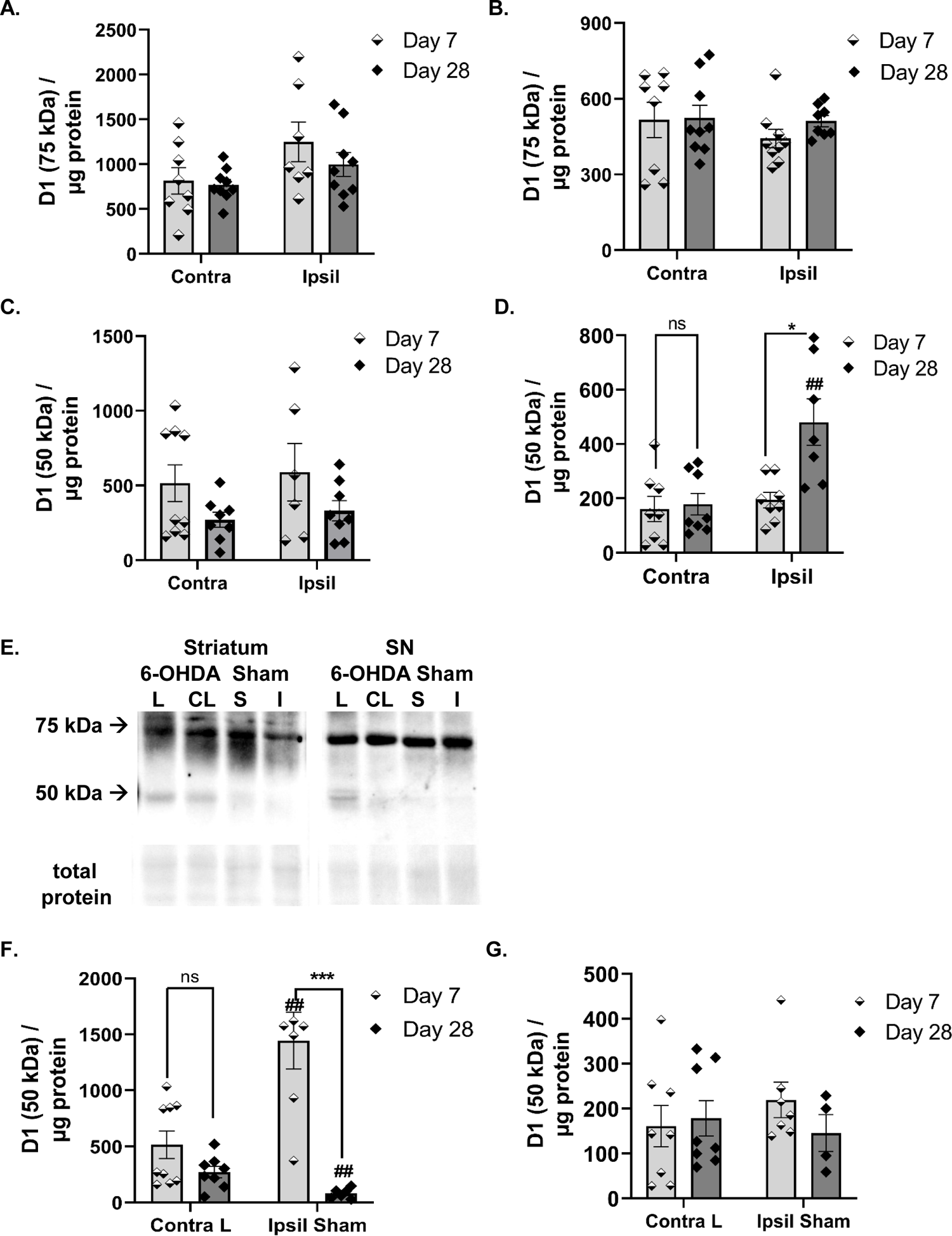
DA D_1_ receptor expression. 6-OHDA group 75 kDa A, B; 50 kDa C,D;. **A. Striatum.** There was an effect of lesion (F_(1,29)_ = 5.4, *p*=0.027), but no effect of time after lesion (F_(1,29)_ = 1.08, ns); or interaction between lesion and time after lesion (F_(1,29)_ = 0.53, ns). No significant differences were seen between any post-hoc comparison. **B. SN.** No effect of lesion (F_(1,14)_ = 1.05, ns), time after lesion (F_(1,16)_ = 0.52, ns), or interaction between lesion and time after lesion (F_(1,14)_ = 0.52, ns). **C. Striatum.** Lesion did not affect expression of the 50 kDa form (F_(1,27)_ = 0.36, ns), but there was an effect on expression time after lesion (F_(1,27)_ = 4.96, *p* =0.035); interaction between lesion and time after lesion (F_(1,27)_ = 0.00, ns). **D. SN.** There was a highly significant effect of lesion (F_(1,12)_ = 22.0, *p* =0.0005), time after lesion (F_(1,15)_ = 5.06, *p* =0.04), and interaction between lesion and time after lesion (F_(1,12)_ = 12.7, *p* =0.004). Day 7 vs day 28; lesioned side (t=2.68, \**p* =0.02, df=14); contralateral to lesioned side (t=0.29, ns, df=14). Contra vs Ipsil to lesion; Day 7 (t=0.90, ns, df=7); Day 28 (t=4.74, ^##^*p* =0.005, df=5). **E. Representative changes in expression of D_1_ receptor, 28 days post lesion.** Differences in expression of the 50kDa form are shown to be restricted in the lesioned SN against other groups and vs side contralateral to lesion. **F. Striatum, contra to lesion.** Sham-operation increased expression of the 50 kDa form vs the side contralateral to lesioned side day 7 after operation. Contra lesion (F_(1,26)_ = 6.62, *p* =0.016); days post-lesion (F_(1,26)_ = 31.4, *p* < 0.0001); contra lesion x days post-lesion (F_(1,26)_ = 15.3, *p =*0.0006). Day 7; contralateral to lesioned side vs Sham (t=3.56, ^##^*p <*0.01, df=14). Day 28; contralateral to lesioned side vs Sham (t=3.11, ^##^*p<*0.01, df=12). Day 7 v day 28; Contralateral to lesion (t=1.75, ns, df=15), Sham (t=4.96, \*\*\**p <*0.001, df=11), **G. SN, contra to lesion.** There was no significant difference in expression of the 50 kDa form between the side contralateral to lesioned side vs. sham-operation. Contra lesion (F_(1,23)_ = 0.08, ns); days post-lesion (F_(1,23)_ = 0.39, ns); contra lesion x days post-lesion (F_(1,23)_ = 1.03, ns).

There was a transient bilateral increase in expression of the 50 kDa form in striatum expression early after lesion (Fig. 9C), likely related to operation itself, since lesion itself did not have a significant effect (Fig. 9F, Suppl. Fig. 12C). A similar increase from sham-operation also occurred in the SN (Fig. 9G, Suppl. Fig. 12D). However, the 6-OHDA lesion increased expression of the 50 kDa form, well-above the increase in the side contralateral to lesion by day 28 after lesion (Fig. 9D). We found no significant difference in expression of the 50 kDa form between the sham-operated side and the side contralateral to lesion (Fig. 9G), signifying that the increase in the 50 kDa form in the SN was lesion-specific and unrelated to sham-operation.

## Discussion

Compensatory mechanisms have been proposed to delay onset of parkinsonian motor signs, including hypokinesia, until TH loss in striatum is substantial [4,7,15–19]. However, the evidence supporting that DA plasticity occurs in striatum in this process is sparse. For example, primates can recover from parkinsonian signs despite severe (∼90%) DA loss, and yet have no difference in striatal DA loss compared with primates with moderate or severe parkinsonian signs [17]. Our study showed that despite virtually complete loss of striatal DA or TH by 7 days, hypokinesia onset was not uniform, and was not decreased overall in two cohorts. Despite the knowledge that DA release in the SN has been established for nearly 50 years [51], consideration of nigral DA signaling in parkinsonian sign onset and severity is largely neglected and therefore sparsely investigated [9, 27-29,37,52–55]. Our systematic and contemporaneous comparison of lesion-induced changes in DA signaling components in striatum and SN revealed the respective roles of DA regulation against the timing of hypokinesia onset. Increased ser31 TH phosphorylation occurred only in the SN, and was sustained, in association with less DA loss against progressive TH protein loss in the lesioned and, later, contralateral to lesioned SN. No consistent changes in ser40 TH phosphorylation occurred in either striatum or SN. Moreover, hypokinesia onset and severity were not uniform after lesion-induction, and nigral, but not striatal, DA loss had coincident correlation against relative decreases in locomotor activity versus prelesion baseline levels. Notably, remaining DA (∼40%) was still more than 3-fold greater than remaining TH protein in the SN by day 28. Despite 60% DA loss in lesioned SN, baseline locomotor activity only decreased overall ∼30%. At that time, DA D_1_ receptor expression increased >2-fold only in lesioned SN. These results point to a two-step compensatory process of increased ser31 TH phosphorylation and, later, increased DA D_1_ receptor levels, exclusively in the SN, that augments DA signaling to mitigate severity of hypokinesia associated with progressive nigrostriatal neuron loss.

Our study addresses long-standing knowledge gaps of how nigrostriatal DA signaling influences movement initiation and maintenance, particularly of internally-guided movement as inferred by open-field activity [56]. Preclinical to clinical studies have shown incongruity between striatal DA signaling and changes in locomotor function [6-14,19,21-29,52-63]. Our study shows similar incongruities; 1) despite >90% loss of TH and DA in striatum 7 days after lesion, locomotor activity did not decrease, and 2) lesion-related increases in D_1_ receptor expression in striatal regions was transient, despite major DA loss. Moreover, there was less DA per remaining TH protein (Fig. 8A) associated with decreased ser31 TH phosphorylation (Fig. 8F), indicating that increased striatal DA turnover did not coincide with increased DA biosynthesis. This argues against increased DA turnover as compensatory mechanism before or after parkinsonian sign onset [7,17,19]. The incongruity between striatal DA function and parkinsonian signs also extends to aging-related parkinsonism. Although hypokinesia also occurs in aging rodent models and humans [9,52,53,57,64–68], no study has reported striatal DA or TH loss exceeding 60% [45,52,57,66–71]. Thus, the outcomes from multiple experimental approaches, from rat to human, indicate that striatal DA deficits are not the sole contributing mechanism of parkinsonian motor deficits, and now extend them by identifying specific mechanisms in the SN to compensate against nigrostriatal neuron loss.

Despite being different compartments of the same neuron, there is disparity in the rate of TH protein loss and DA regulation between striatum and SN during nigrostriatal neuron loss. These disparities emulate human PD, wherein TH loss also proceeds at a much faster rate in striatal regions [6,58,60]. Therefore, because DA loss is mitigated in SN, but not in striatum, this autonomous regulation points to the possibility that DA regulation in the SN alone influences hypokinesia onset [7,27,28,62] or its alleviation [9,10, 52]. Thus, although DA release occurs in both areas during nigrostriatal neuron activity [73–75], its impact on locomotor function may be differentially-influenced by striatum and SN. In this study, compensatory processes to maintain DA signaling against TH protein, eventual nigrostriatal cell and DA loss were engaged solely in the SN. These results beg consideration of a paradigm shift to fully understand the role of DA in parkinsonian motor sign etiology and treatment..

Tyrosine hydroxylase phosphorylation is a primary post-translational modification of activity [45, 76] and the dichotomy in ser31 phosphorylation stoichiometry between SN and striatum is well-established [9,31,32,45–47,62]. However, it is not known how site-specific TH phosphorylation may alter DA levels during progressive nigrostriatal neuron loss [31–34]. Phosphorylation at ser40 has been long-assumed to be the primary post-translational modification to regulate L-DOPA biosynthesis [76], but *in vivo* evidence to substantiate this mechanism is tenuous [31,45–47,62]. Nigrostriatal neuron activity drives DA release needed for initiation of locomotor activity [73–75]. Thus, increased TH phosphorylation would replenish released DA, which would be even more critical during TH protein and cell loss in PD. A 1.6-fold increase ser31 phosphorylation from baseline levels of 0.08 increases L-DOPA biosynthesis *in situ* [77]. Similar ranges of stoichiometry at ser31 in the SN are seen in this study. Moreover, multiple *in vivo* studies show congruency between differences in ser31 TH phosphorylation stoichiometry and DA tissue content [31,45–47,62]. Importantly levels of ser40 phosphorylation stoichiometry *in situ* are also comparable to *in vivo* studies [9,31,45–47,62,78]. However, the levels of ser40 *in vivo* may not be sufficient to affect L-DOPA biosynthesis [31,45,47,62,78], including in striatum [35, 78], as even two-fold increases (from 0.03 to 0.06) do not increase L-DOPA biosynthesis [77]. Moreover, increased ser40 phosphorylation may promote proteosomal degradation of TH [31,79,80], which may explain accelerated TH loss in the human striatum of PD patients [35]. We speculate the mechanism driving increased ser31 is related in part to diminished DA-mediated stimulation of DA D_2_ autoreceptors [78]. Thus, ser31 TH phosphorylation may increase in the SN from the lack of extracellular DA caused by lesion, although this possibility is beyond the scope of our current study and remains to be investigated.

Late after lesion, DA D_1_ receptor expression increased in lesioned SN, when DA loss reached∼60%. Curiously, despite early and sustained severe DA loss in striatum, there was only transient and modest increase in DA D_1_ expression. Thus, these two events suggest that DA D_1_ expression may increase in response to decreased DA only in SN, and likely has functional significance as DA D_1_ expression or function in SN alone may affect locomotor function [9,29,53]. DA D_1_ receptors in the SN are on striatonigral terminals [41, 43], and originate from striatum, as mRNA expression for D_1_ is not seen in SN [41, 81]. Activation of D_1_ receptors in the SN increases GABA release [39, 50], which mitigates the inhibitory output of the basal ganglia to promote movement initiation [37]. The critical role of D_1_ function in movement initiation in PD is not as well-elucidated compared to the DA D_2_ receptor [82]. However, studies in parkinsonian primates and patients alike show D_1_ receptor agonists reduce parkinsonian severity [36, 83]. This locomotor-promoting effect may be related to the upregulation of receptor expression observed here, and augment striatonigral GABA release [84]. To this point., despite 60% loss of nigral DA at the late stage of lesion, there was only ∼25% decreased locomotor activity. As a point of reference, experimentally-induced decreases in DA ∼40% in the SN following nigral TH inhibition, reduce locomotor activity by a similar magnitude (∼35%) in young rats [28]. Therefore, given even greater DA loss of 60% by day 28, it is plausible increased DA D_1_ receptor expression in the SN offset DA loss to mitigate hypokinesia severity.

The neural mechanisms of PD involve loss of nigrostriatal DA neurons and pathology related to a-synuclein and/or Lewy body neuronal processes [3]. In human PD, >80% TH loss in striatum has occurred at the time of motor impairment, and greater than in SN [6, 7]. Our study showed no correlation between the degree of TH loss between striatum and SN, indicating different processes govern TH loss between these regions, as suggested in human studies [22,58,87,88].. In human PSD, motor decline worsens despite virtually complete loss of striatal DA markers [6,58–60] within 5 years diagnosis [6], and our results show a similar dichotomy. Yet, motor impairments are still framed in context of striatal DA loss, and measures in the SN, if any, are largely confined to gauging neuronal loss. Our study also shows DA loss paralleled severe TH loss in striatum, whereas in the SN, DA loss was much less and correlative with the magnitude of hypokinesia severity. These observations may be applicable to detecting PD in the prodromal period, wherein non-motor impairments, including specific executive function deficits [85] are underway during this period [1–4]. To this end, there was major DA loss in the nucleus accumbens, contrasting against increased DA in the VTA 7 days after lesion induction. There was also eventual loss of TH protein in the VTA, albeit at a slower rate than in SN, which also occurs in human PD. There also was an eventual loss of TH protein in the SN contralateral to lesion that was not seen in striatum, which is also reported in other rat PD models [26, 86]. This change suggests that the progression from Hoehn Yahr Stage 1 to 2, may be in part related to this nigra-specific change. Thus, the mechanisms that increase nigral DA signaling to confer resiliency against TH protein and neuronal loss in SN may play a role in the heterogeneity of the timing of hypokinesia-specific deficits [22,23,59,60,87–91].

Our results pose new questions. First, why does forelimb use diminish, but not locomotor activity in the early stages of TH loss in the SN, even though diminished forelimb use was essential for eventual decline in locomotor activity. We do not know is compensatory mechanisms augmented DA signaling in striatum earlier than 7 days post-lesion and if forelimb use could be partially recovered if striatal DA loss was mitigated. We also observed increased forelimb use in the intact side by ∼20%, which may not only translate to less decline in the open-field but also signal that DA release dynamics increase contralateral to lesion, despite no DA loss in either striatum or SN between 7 and 28 days after lesion, as suggested by work in primate models [90].

### Summary

Our results provide translation to human PD and new mechanistic insight. First, imaging studies in PD patients should interrogate DA signaling status in the SN as another measure of PD progression and motor symptom severity. To this point, despite major loss of DA tissue content, TH protein in striatum, hypokinesia was not evident after nigrostriatal lesion. Moreover, eventual decline of DA content only in the SN was correlated with decreased locomotor activity. Increased ser31 TH phosphorylation offset DA loss against TH protein loss and gradual neuron loss only in the SN, arguing that increased DA signaling in the SN is critical for locomotor activity, as suggested in other studies [9,29,52,53,61,62]. Moreover, increased DA D_1_ receptors countered loss of DA and likely mitigated the severity of parkinsonian signs. The results have revealed that the plasticity in DA signaling in remaining nigrostriatal neurons occurs in the SN and delays hypokinesia onset and severity. Accordingly, our study address a long-standing knowledge gap in why parkinsonian sign onset occurs only after substantial loss of DA proteins in the striatum. A new and sustained focus on DA regulation in the SN can further address the molecular origins of vulnerability and resiliency to hypokinesia, as well as therapeutic avenues to treat parkinsonian signs.

## Acknowledgements

Funding: This study was funded by the US Department of Defense U.S. Army Medical Research and Material Command Congressionally Directed Medical Research Program (Award Number W81XWH-19-1-0757) to MFS and Office of Vice president for Research and Innovation at University of North Texas Health Science Center and Visiting Scholar Fellowship from the Parkinson’s Foundation to EAK. Additional support was provided by R01ES033892 and U01NS108956 to JRR. and a CV Starr Fellowship to AB.

