## Supplemental Data for "Nigral-specific increase in ser31 tyrosine hydroxylase phosphorylation offsets dopamine loss and forestalls hypokinesia onset during progressive nigrostriatal neuron loss"

**Supplemental results**

**Figure S1.**

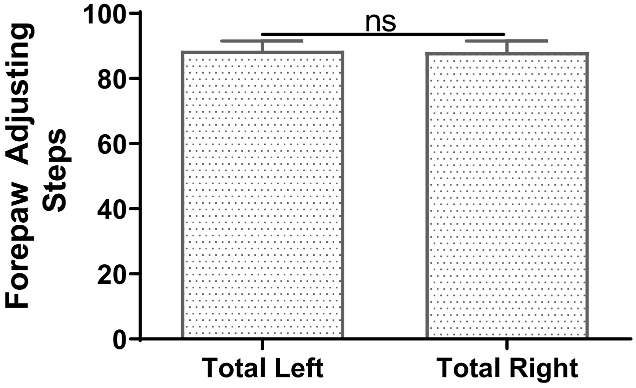


**Suppl. Figure 1. Baseline FAS performance in cohort prior to Sham operation or 6-OHDA lesion.** Left forelimb use was not significantly different from right forelimb use in rats prior to assignment into sham operation of the medial forebrain bundle or nigrostriatal 6-OHDA lesion ($t = 0.24$, ns, $df = 18$).

**Figure S2.**

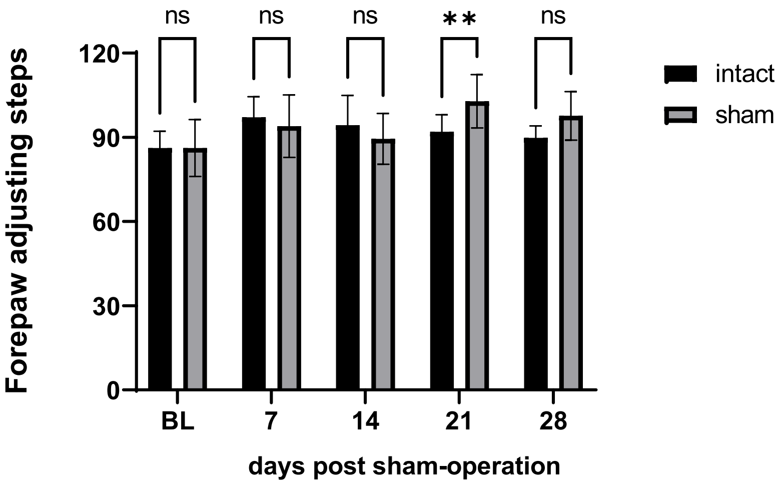


**Suppl. Figure 2. Sham-operation impact on FAS.** Sham-operation on the medial forebrain bundle did not have a significant effect on forelimb use against forelimb use associated with the intact side ($F_{(1,5)}$ = 0.81, ns). However, forelimb use increased over time post-surgery ($F_{(4,20)}$ = 4.15, *p*=0.01) and significant interaction with sham-operation X time post-surgery ($F_{(4,20)}$ = 5.83, *p*<0.003). Day 21 (t=3.80, ***p*=0.006, Bonferroni Multiple Comparison Test).

Figure S3.

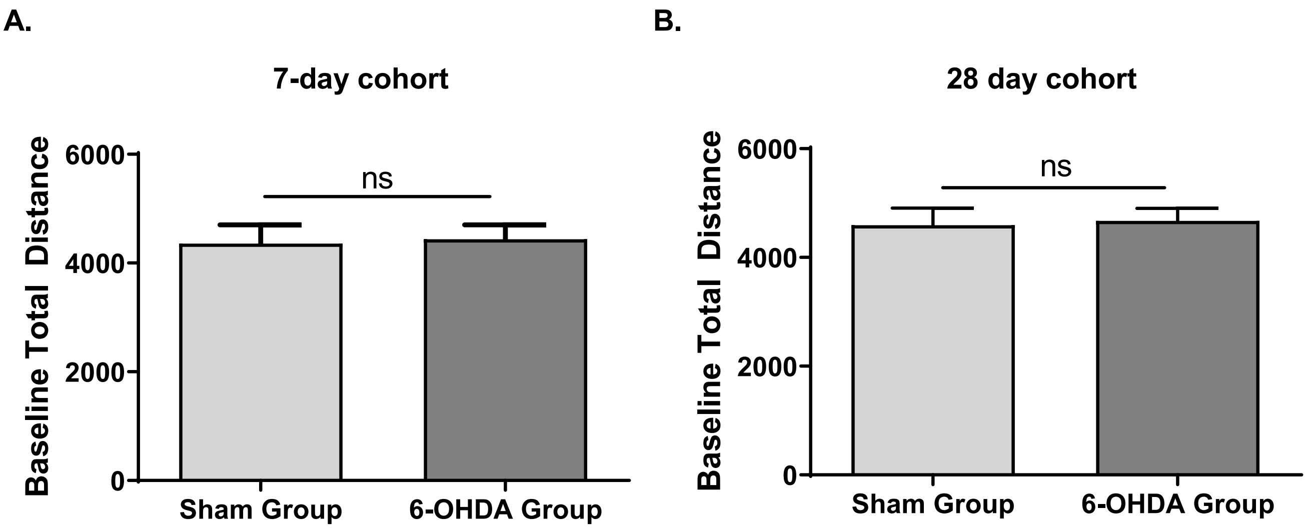


**Suppl. Figure S3. Equivalent baseline locomotor activity between cohorts and prior to assignment into treatment groups. A. 7-day cohort.** Total distance was not significantly different between rats assigned to the sham or 6-OHDA group (t=0.18, ns, df=18). **B. 28-day cohort.** Total distance was not significantly different between rats assigned to the sham or 6-OHDA group (t=0.202, ns, df=18).

Figure S4.

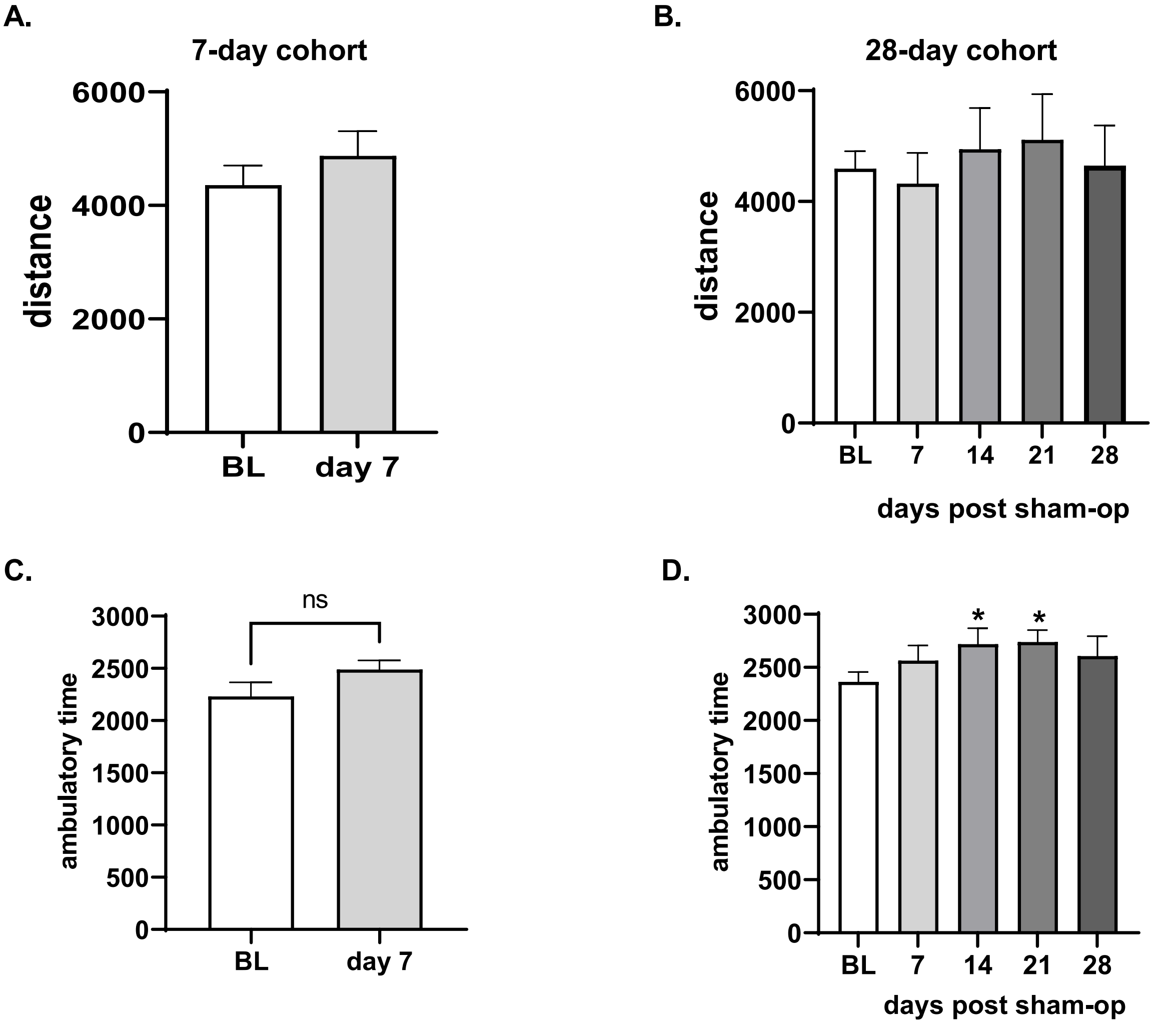


**Suppl. Figure S4. Sham-operation had minimal impact on locomotor activity. A. Total distance, 7-day cohort.** Total distance was not significantly different at day 7 (t=1.91, ns, df=7). **B. Total distance, 28-day cohort.** Total distance was not significantly different across all days evaluated ($F_{(4,24)}$ = 0.88, ns). **C. Ambulatory time, 7-day cohort.** Time spent moving was not significantly different at day 7 (t=1.90, ns, df=7). **D, Ambulatory time, 28-day cohort.** Time spent moving increased ~15% above baseline (BL) at two time points (day 14, 21) across all days evaluated ($F_{(4,24)}$ = 3.29, *p* =0.03); day 7 (q=1.71, ns), day 14 (q=3.03, *p*<0.05), day 21 (q=3.20, *p*<0.05), day 28 (q=2.07, ns), Dunnett's multiple comparisons test.

**Figure S5.**

**A.**

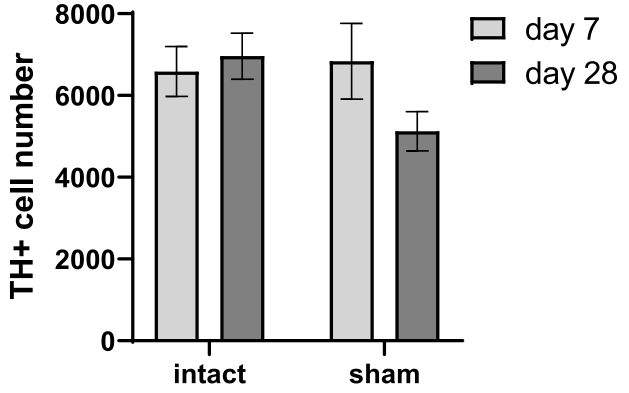


**B.**

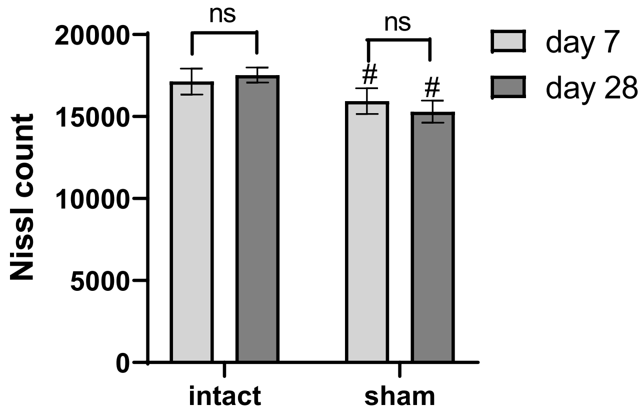


**Suppl. Figure S5. Minimal impact of sham-operation on TH cell number or total cell number (Nissl) in SN. A. TH cell number.** Sham-operation ($F_{(1,7)}$ = 3.72, ns (*p*=0.10)); days post-sham-operation ($F_{(1,8)}$ = 0.46, ns); sham-operation x days post sham-operation ($F_{(1,7)}$ = 6.12, $p$ <0.05). Day 7 vs day 28; intact side (t=0.44, ns); sham-op side (t=1.64, ns); Intact vs. sham-op side; Day 7 (t=0.60, ns); Day 28 (t=2.40, ns). **B. Total cell number (Nissl).** Sham-operation ($F_{(1,9)}$ = 16.76, *p*=0.003); days post-sham-operation ($F_{(1,9)}$ = 0.02, ns); sham-operation x days post sham-operation ($F_{(1,9)}$ = 1.54, ns). Day 7 vs day 28; intact side (t=0.41, ns); sham-op side (t=0.61, ns); Intact vs. sham-op side; Day 7 (t=2.74, $^{\#}p$ <0.05); Day 28 (t=2.94, $^{\#}p$ <0.05).

**Figure S6.**

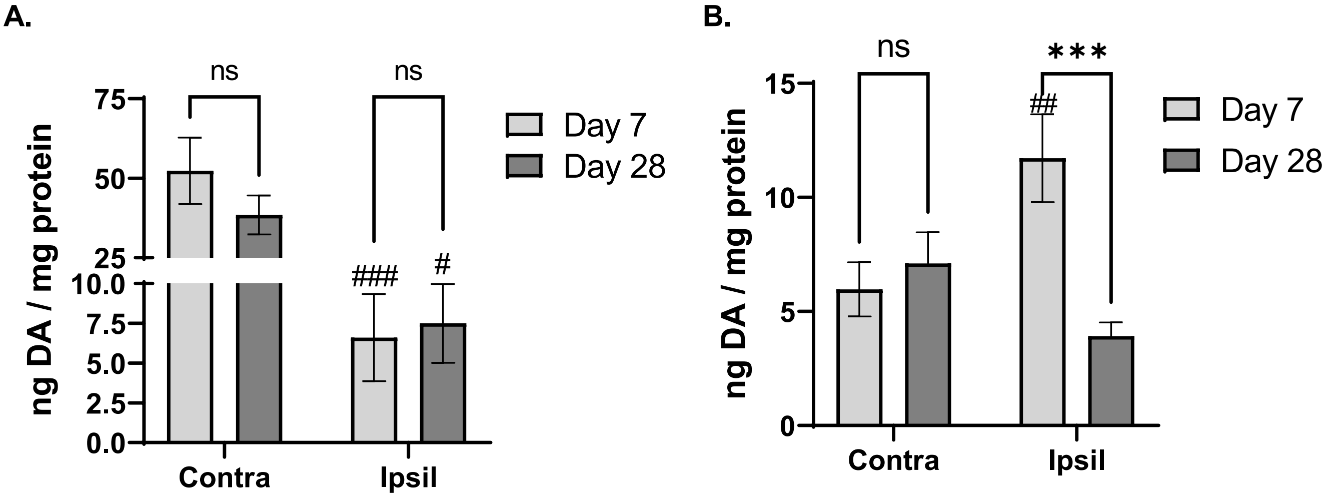


**Suppl. Figure S6. 6-OHDA lesion impact on DA tissue content in mesoaccumbens pathway. A. NAc.** Lesion ($F_{(1,16)}$ = 31.6, $p$<0.0001); days post-lesion ($F_{(1,16)}$ = 1.26, ns); lesion x days post-lesion ($F_{(1,15)}$ = 1.17, n). Day 7 vs day 28; lesioned side (t=0.10, ns); contralateral to lesioned side (t=1.16, ns). Contra vs Ipsil to lesion; Day 7 (t=4.74, $^{###}p$ <0.001); Day 28 (t=3.21, $^{#}p$ <0.05). **B. VTA.** Lesion ($F_{(1,15)}$ = 1.47, ns); days post-lesion ($F_{(1,16)}$ = 4.88, $p$=0.042); lesion x days post-lesion ($F_{(1,15)}$ = 17.61, $p$<0.001). Day 7 vs day 28; lesioned side (t=4.17, ***$p$ <0.001); contralateral to lesioned side (t=0.62, ns). Contra vs Ipsil to lesion; Day 7 (t=3.75, $^{##}p$ <0.01); Day 28 (t=2.15, ns).

**Figure S7.**

**A.**

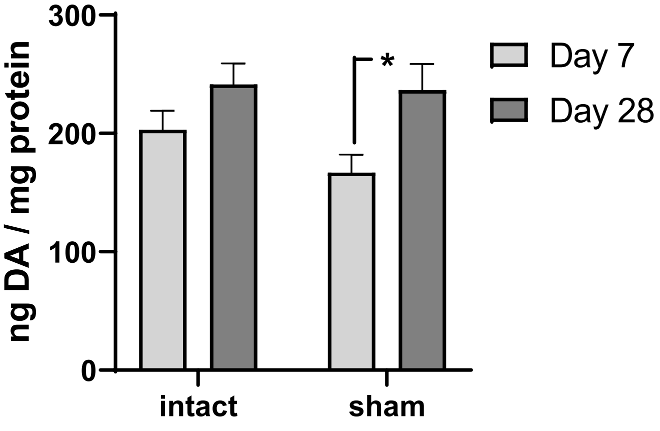


**B.**

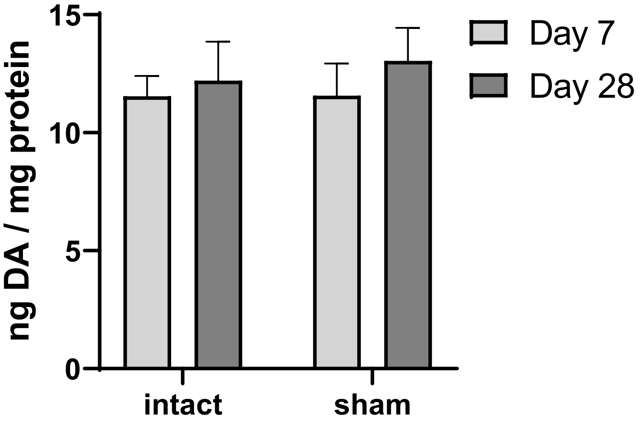


**C.**

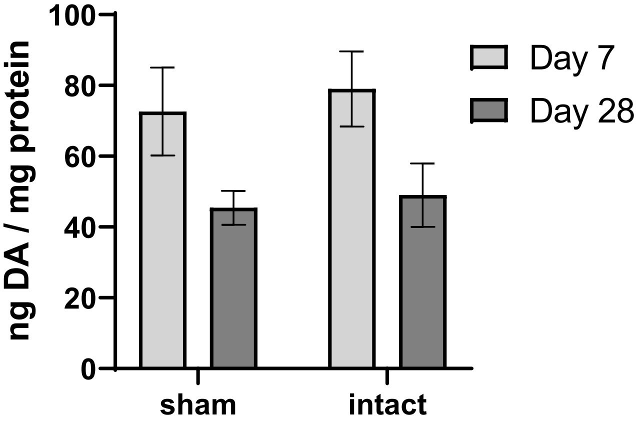


**D.**

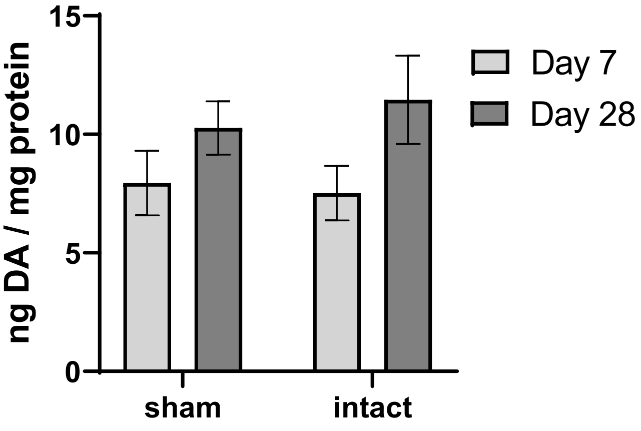


**Suppl. Figure S7. Sham-operation impact on DA tissue content. A. Striatum.** Sham-operation ($F_{(1,13)}$ = 2.15, ns); days post-sham-operation ($F_{(1,13)}$ = 6.74, *p*=0.022); sham-operation x days post sham-operation ($F_{(1,13)}$ = 1.28, ns). Sham-op side, Day 7 vs day 28 (t=2.79, $^{\#}p$ =0.02). **B. SN.** Sham-operation ($F_{(1,13)}$ = 0.11, ns); days post-sham-operation ($F_{(1,13)}$ = 0.61, ns); sham-operation x days post sham-operation ($F_{(1,13)}$ = 0.10, ns). **C. NAc.** Sham-operation ($F_{(1,13)}$ = 0.670, ns); days post-sham-operation ($F_{(1,13)}$ = 5.07, *p* =0.04); sham-operation x days post sham-operation ($F_{(1,13)}$ = 0.05, ns). **D. VTA.** Sham-operation ($F_{(1,13)}$ = 0.22, ns); days post-sham-operation ($F_{(1,13)}$ = 3.04, ns); sham-operation x days post sham-operation ($F_{(1,13)}$ = 0.98, ns).

**Figure S8.**

**A.** **B.**

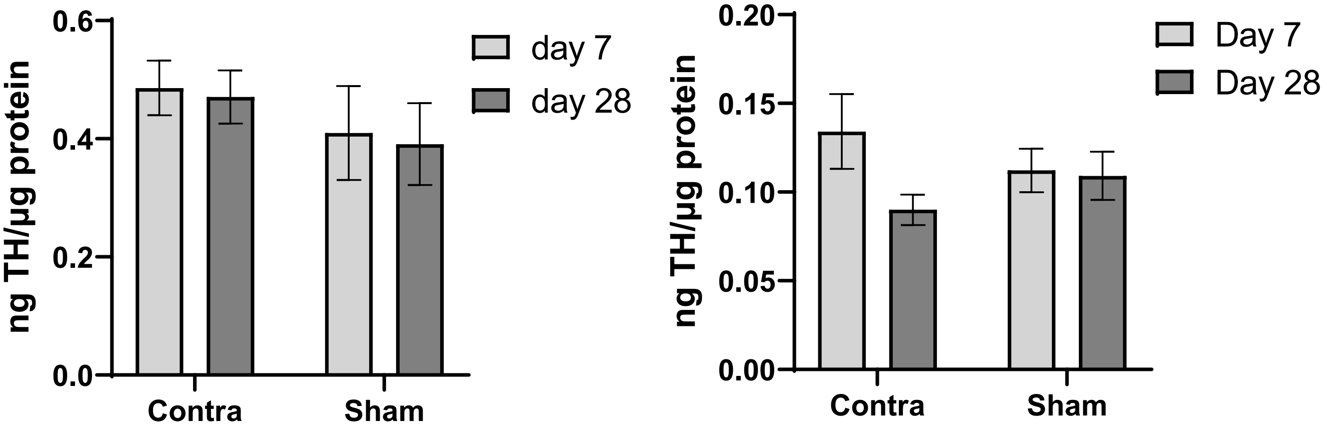


**Suppl. Figure S8. Sham-operation had no effect on TH protein expression. A. Striatum.** Sham-operation did not affect TH protein levels in striatum. Sham-op ($F_{(1,12)}$ = 2.80, ns); days post-sham-operation ($F_{(1,13)}$ = 0.04, ns); sham-operation x days post sham-operation ($F_{(1,12)}$ = 0.00, ns). **B. SN.** Sham-operation did not affect TH protein levels in SN. Sham-op ($F_{(1,10)}$ = 0.001, ns); days post-sham-operation ($F_{(1,12)}$ = 2.10, ns); sham-operation x days post sham-operation ($F_{(1,10)}$ = 2.31, ns).

Figure S9

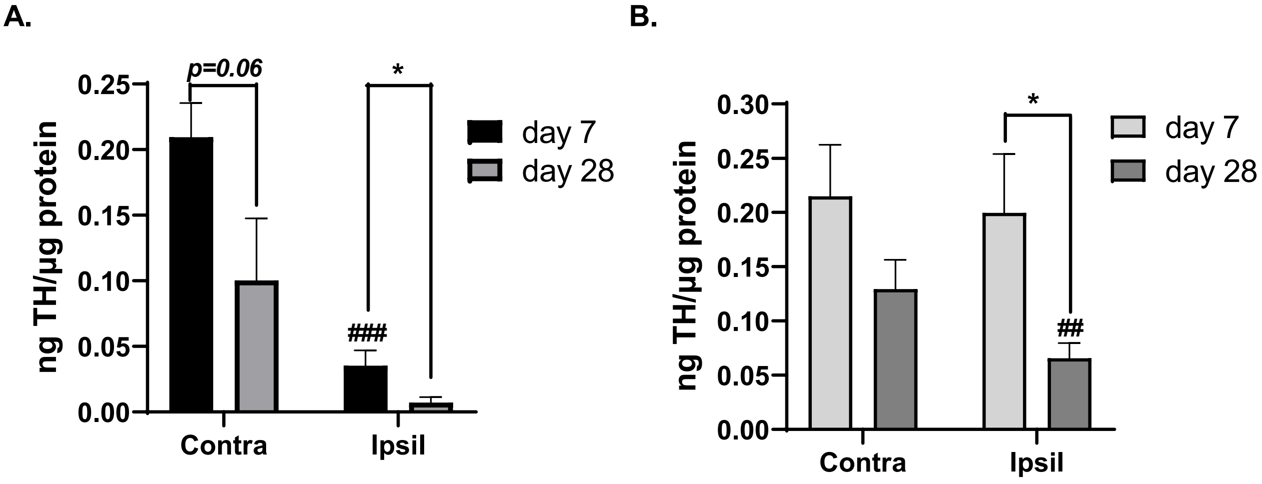


**Suppl. Figure S9. 6-OHDA lesion impact on TH protein expression in mesoaccumbens pathway. A. NAc.** Lesion ($F_{(1,16)}$ = 21.7, *p*=0.0003); days post-lesion ($F_{(1,16)}$ = 6.62, *p* =0.02); lesion x days post-lesion ($F_{(1,16)}$ = 1.99, ns). Day 7 vs day 28; lesioned side (t=2.56, **p* <0.05); contralateral to lesioned side (t=2.02, *p* =0.06). Contra vs Ipsil to lesion; Day 7 (t=5.32, ###*p* =0.0007); Day 28 (t=1.69, ns). **B. VTA.** Lesion ($F_{(1,16)}$ = 3.82, *p* =0.07); days post-lesion ($F_{(1,16)}$ = 5.22, *p*=0.036); lesion x days post-lesion ($F_{(1,15)}$ = 2.54, ns). Day 7 vs day 28; lesioned side (t=2.54, **p* <0.05); contralateral to lesioned side (t=1.57, ns). Contra vs Ipsil to lesion; Day 7 (t=0.12, ns); Day 28 (t=3.88, ##*p* <0.01).

**Figure S10.**

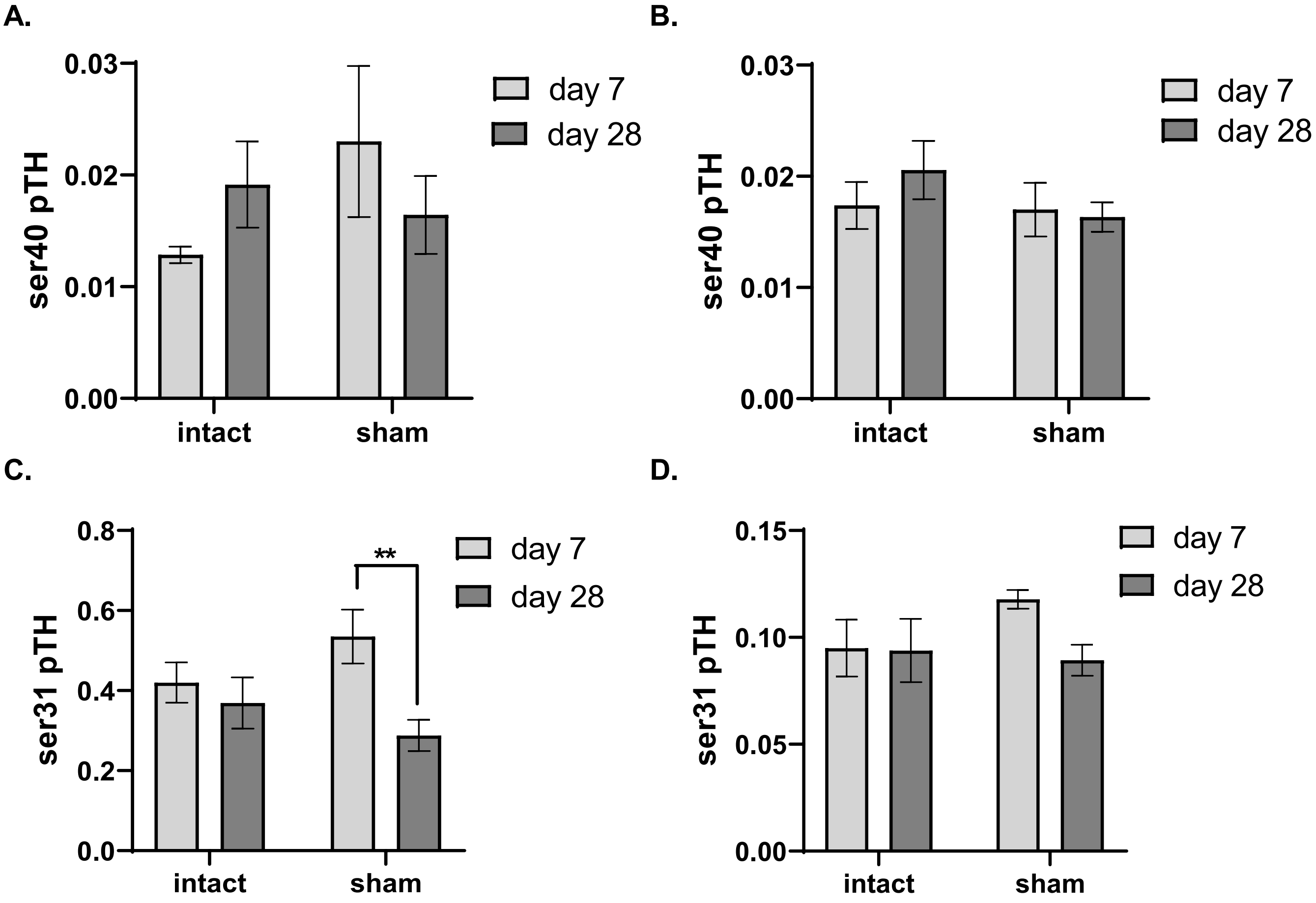


**Suppl. Figure S10. TH phosphorylation stoichiometry (PS) in sham-op group. A. Striatum, ser40 PS** Sham-operation ($F_{(1,11)}$ = 1.09, ns); days post-sham-operation ($F_{(1,13)}$ = 0.00, ns); sham-operation x days post sham-operation ($F_{(1,11)}$ = 3.23, *p* =0.10). **B. SN, ser40 PS** Sham-operation ($F_{(1,12)}$ = 1.36, ns); days post-sham-operation ($F_{(1,13)}$ = 0.32, ns); sham-operation x days post sham-operation ($F_{(1,12)}$ = 0.91, ns). **C. Striatum, ser31 PS** Sham-operation ($F_{(1,11)}$ = 0.59, ns); days post-sham-operation ($F_{(1,13)}$ = 4.29, ns); sham-operation x days post sham-operation ($F_{(1,11)}$ = 11.63, *p*=0.006). Sham-op side, Day 7 vs day 28 (t=3.17, ***p* =0.008). **D. SN, ser31 PS** Sham-operation ($F_{(1,11)}$ = 0.75, ns); days post-sham-operation ($F_{(1,13)}$ = 0.58, ns); sham-operation x days post sham-operation ($F_{(1,11)}$ = 0.63, ns).

**Figure S11.**

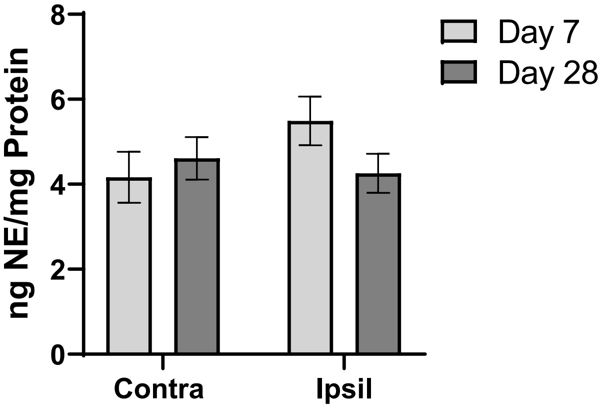


**Suppl. Figure S11. Norepinephrine levels in SN were unaffected by 6-OHDA lesion.**

Lesion ($F_{(1,12)}$ = 0.93, ns); days post-lesion ($F_{(1,13)}$ = 0.48, ns); lesion x days post-lesion ($F_{(1,12)}$ = 2.80, ns).

**Figure S12.**

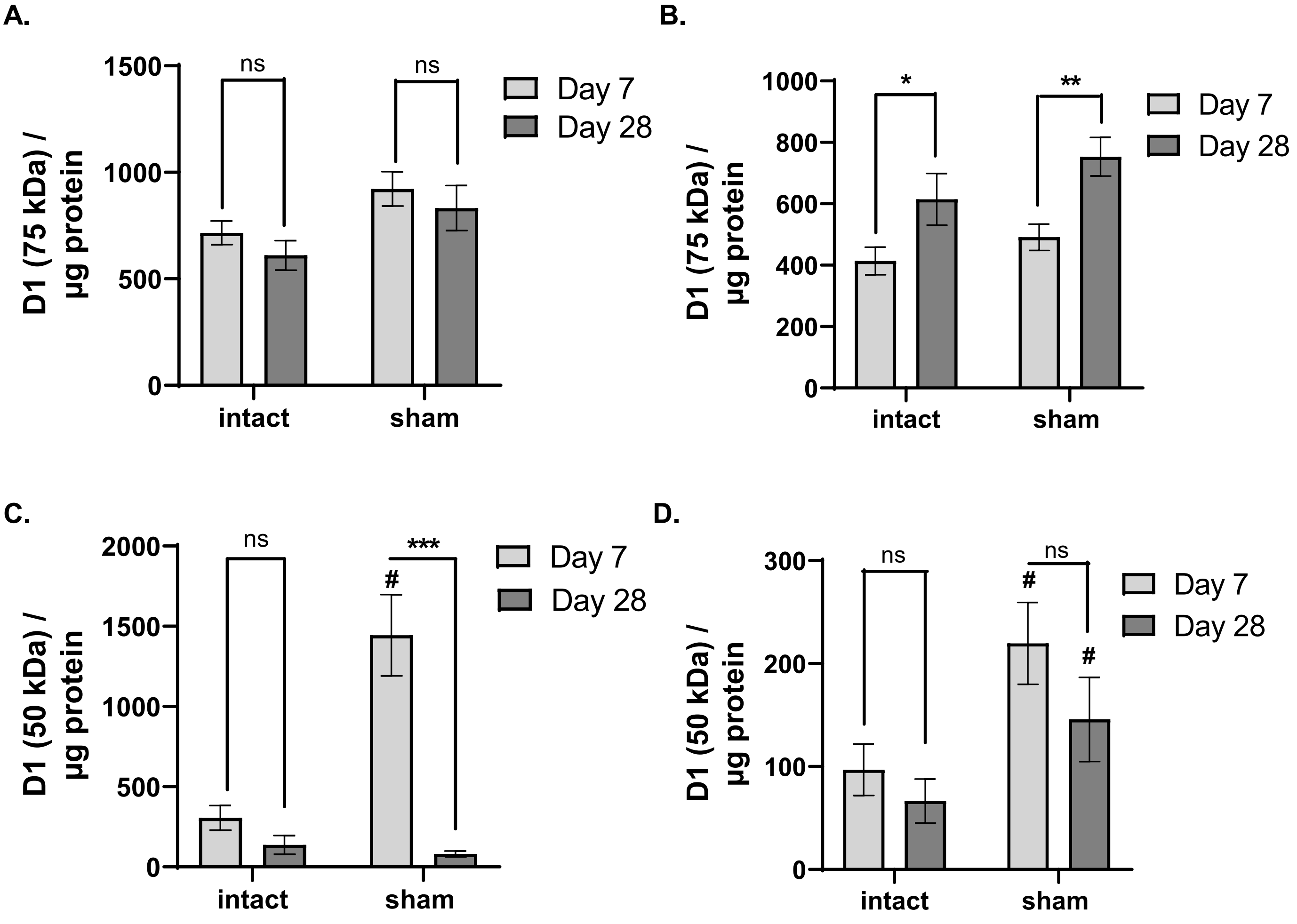


**Suppl. Figure S12. DA $D_1$ receptor expression in sham-operation. A. Striatum, 75 kDa.** Sham-operation ($F_{(1,11)}$ = 7.24, $p$ =0.021); days post-sham-operation ($F_{(1,13)}$ = 1.49, ns); sham-operation x days post sham-operation ($F_{(1,11)}$ = 0.01, ns). **B. SN, 75 kDa.** Sham-operation increased $D_1$ receptor expression ($F_{(1,10)}$ = 5.3, $p$=0.04); days post-sham-operation ($F_{(1,12)}$ = 11.9, $p$=0.005); sham-operation x days post sham-operation ($F_{(1,10)}$ = 0.29, ns). Day 7 vs day 28; intact side (t=2.20, *$p$=0.05); sham-op side (t=3.540, **$p$ =0.005); Intact vs. sham-op side; Day 7 (t=1.68, ns); Day 28 (t=1.77, ns). **C. Striatum, 50 kDa.** Sham-operation ($F_{(1,9)}$ = 13.7, $p$=0.005); days post-sham-operation ($F_{(1,13)}$ = 27.3, $p$=0.0002); sham-operation x days post sham-operation ($F_{(1,9)}$ = 16.8, $p$ =0.003). Day 7 vs day 28; intact side (t=1.69, ns); sham-op side (t=4.96, ***$p$ =0.0004); Intact vs. sham-op side; Day 7 (t=3.66, $^{\#}p$ <0.05); Day 28 (t=0.86, ns).

**D. SN, 50 kDa.** Sham-operation increased $D_1$ receptor expression ($F_{(1,9)}$ = 11.1, $p$=0.009); days post-sham-operation ($F_{(1,9)}$ = 1.63, ns); sham-operation x days post sham-operation ($F_{(1,9)}$ = 0.51, ns). Day 7 vs day 28; intact side (t=0.81, ns); sham-op side (t=1.20, ns); Intact vs. sham-op side; Day 7 (t=2.86, $^{\#}p$ <0.05); Day 28 (t=3.36, $^{\#}p$ <0.05).
